# Activation of Neurotensin Receptor 1 in entorhinal cortex rescues olfactory generalization in two mouse models of autism

**DOI:** 10.64898/2026.07.29.741644

**Authors:** Kassandra L. Sturm, Daryna Semak, Mitchell Swerdloff, Bardia Askari, Raddy L. Ramos, Gonzalo H. Otazu

**Affiliations:** New York Institute of Technology, College of Osteopathic Medicine

## Abstract

Individuals with Autism Spectrum Disorder (ASD) show impaired transfer of learned information to novel contexts ^1^, a computational deficit proposed to contribute to atypical sensory processing in ASD ^2^. Mouse models carrying mutations in the ASD-associated genes *Shank3*^3^ and *Cntnap2*^4^ show preserved target odor identification in familiar backgrounds but impaired generalization to novel odor backgrounds compared with wild-type mice^5,6^. Olfactory circuits are disrupted in these models at multiple levels, including the olfactory bulb^7,8^ and downstream areas such as the cortical amygdala^9^ and prefrontal cortex^10^. However, it remains unknown whether restoring olfactory circuit function can rescue impaired sensory generalization.

Here, we show that pharmacological activation of neurotensin receptor 1 (Ntsr1) restores the ability of both *Cntnap2^−/−^*and *Shank3B^+/−^* mice to recognize learned target odors in novel odor backgrounds. We further identify the entorhinal cortex as the principal site mediating this effect. Activation of Ntsr1 in the entorhinal cortex reduced olfactory bulb responses to novel background odors, increasing the similarity between neural representations of novel and previously learned odor mixtures. These changes improved neural generalization and restored behavioral recognition of target odors in novel sensory environments. Together, these results identify the entorhinal cortex as a top-down regulator of early olfactory processing that can rescue sensory generalization deficits in mouse models of ASD. Our findings establish pharmacological Ntsr1 activation in entorhinal cortex as a novel therapeutic target for sensory dysfunction in ASD.

## Main

Individuals with Autism Spectrum Disorder (ASD) exhibit a strong preference for routine and predictable sensory experiences, with novel stimuli often eliciting distress. This clinical feature is captured in the DSM-5 as “insistence on sameness”^11^. Consistent with this observation, computational accounts have proposed that ASD is associated with deficits in generalizing learned information to novel contexts^1^. Although these cognitive and sensory difficulties substantially affect quality of life, there are currently no FDA-approved pharmacological treatments that specifically address them. Identifying neural circuits that can be modulated through systemic pharmacological interventions may therefore provide new therapeutic opportunities for reducing atypical responses to novelty in ASD.

Consistent with clinical observations in ASD^12^, mouse models carrying mutations in ASD-associated genes exhibit atypical responses to novel odors^13,14^. *Shank3B^+/−^*^3^ and *Cntnap2^−/−^* ^4^ mice are impaired in their ability to generalize learned target odors to novel odor backgrounds^5,6^. This deficit parallels computational accounts of ASD that emphasize impaired generalization to novel contexts^2^.

Circuit deficits in olfactory processing in mouse models carrying ASD-associated mutations can be detected as early as the output of the olfactory bulb. Olfactory receptor neurons (ORNs) in the olfactory epithelium project to specific glomeruli in the olfactory bulb, where sensory input is transformed by local inhibitory circuits and relayed to downstream brain regions by the principal output neurons, mitral and tufted (M/T) cells^15–17^. M/T cells in *Fmr1^−/−^*mice exhibit enhanced sensitivity to glomerular activation^13^. In addition, extensive training with a background odor improves the ability of *Cntnap2^−/−^* and *Shank3B^+/−^*mice to detect a familiar target odor in that background^5,6^, and this behavioral improvement is accompanied by a reduction in odor-evoked activity in olfactory bulb M/T cells^8^. Together, these findings suggest that attenuating odor-evoked olfactory bulb output may improve odor generalization in mouse models carrying ASD-associated mutations.

M/T cell activity can be attenuated through genetic, optogenetic, and pharmacological manipulations. Activity can be reduced either by decreasing excitatory input to M/T cells^18^ or by activating inhibitory interneuron populations within the olfactory bulb^19–21^. Inhibitory interneurons in the OB that act on M/T cells can also be modulated indirectly through descending feedback projections originating from the piriform cortex^22,23^ and the anterior olfactory nucleus (AON) ^24–27^.

Although effective experimentally, optogenetic and locally administered pharmacological manipulations are inherently invasive and difficult to translate into therapeutics for ASD. We therefore sought to identify endogenous receptors that could be targeted by systemically administered drugs to selectively engage neural populations capable of suppressing olfactory bulb output. Several manipulations that reduce M/T cell activity act through GABA_B_ receptors ^18^ and oxytocin receptors^25^, making them amenable to pharmacological intervention. However, these receptors have already been targeted in clinical trials for ASD using arbaclofen^28^ (Veenstra-VanderWeele et al., 2017) and intranasal oxytocin^29^, respectively.

We focused on Neurotensin Receptor 1 (Ntsr1) because it has not previously been targeted in clinical trials for ASD. Optogenetic activation of Ntsr1-expressing neurons in the piriform cortex using the Ntsr1-Cre GN209 mouse line generated by the GENSAT project^30^ recruits descending projections to the olfactory bulb^22^. We hypothesized that systemic administration of an Ntsr1 agonist could also preferentially recruit Ntsr1-positive piriform cortex neurons that project to the olfactory bulb. This strategy is supported by the low levels of neurotensin binding in the adult brain ^31^, which may permit selective pharmacological activation of Ntsr1-expressing neurons in piriform cortex. To test this hypothesis, we examined whether systemic activation of Ntsr1 could improve odor generalization in mouse models carrying ASD-associated mutations.

Ntsr1 is a G-protein-coupled receptor^32^ that has been implicated in the regulation of feeding behavior and dopaminergic signaling^33^. Ntsr1 is the highest-affinity receptor for neurotensin, a 13-amino-acid neuropeptide^34^ that regulates diverse physiological processes including blood pressure, body temperature, pain perception, and feeding^35^. Native neurotensin, however, is poorly suited for pharmacological manipulation because it is rapidly inactivated by peptidases in vivo^36,37^. To overcome this limitation, we used PD149163, a metabolically stable neurotensin analog that is resistant to peptidase degradation and readily crosses the blood–brain barrier^38^. PD149163 binds selectively to Ntsr1 with negligible affinity for other neurotensin receptors or unrelated receptor systems, including dopaminergic and serotonergic receptors^39^. Its favorable pharmacokinetic properties and brain penetration have led to its evaluation as a systemically administered therapeutic agent in preclinical models of schizophrenia^39–42^. We therefore used PD149163 as a pharmacological tool to selectively activate Ntsr1-expressing neurons using subcutaneous injections.

### Subcutaneous injections of Neurotensin Receptor 1 Agonist rescued target recognition in novel background odors in *Shank3B^+/−^ and Cntnap2^−/−^* mice

To determine whether activation of Ntsr1 could improve generalization to novel sensory contexts, we used an olfactory CAPTCHA task in which head-fixed mice identified a familiar target odor against a novel background odor^5^ (see **Figure 1A-C**) Previous studies have shown that *Cntnap2*^−/−^ mice ^4^ (JAX Stock No. 017482) and *Shank3B*^+/−^ mice^3^ (JAX Stock No. 017688) are impaired in this task relative to wild-type controls when the background odor is novel, but performed similarly to WT mice when tested with background odors encountered during training ^5,6^.

**Figure 1.**
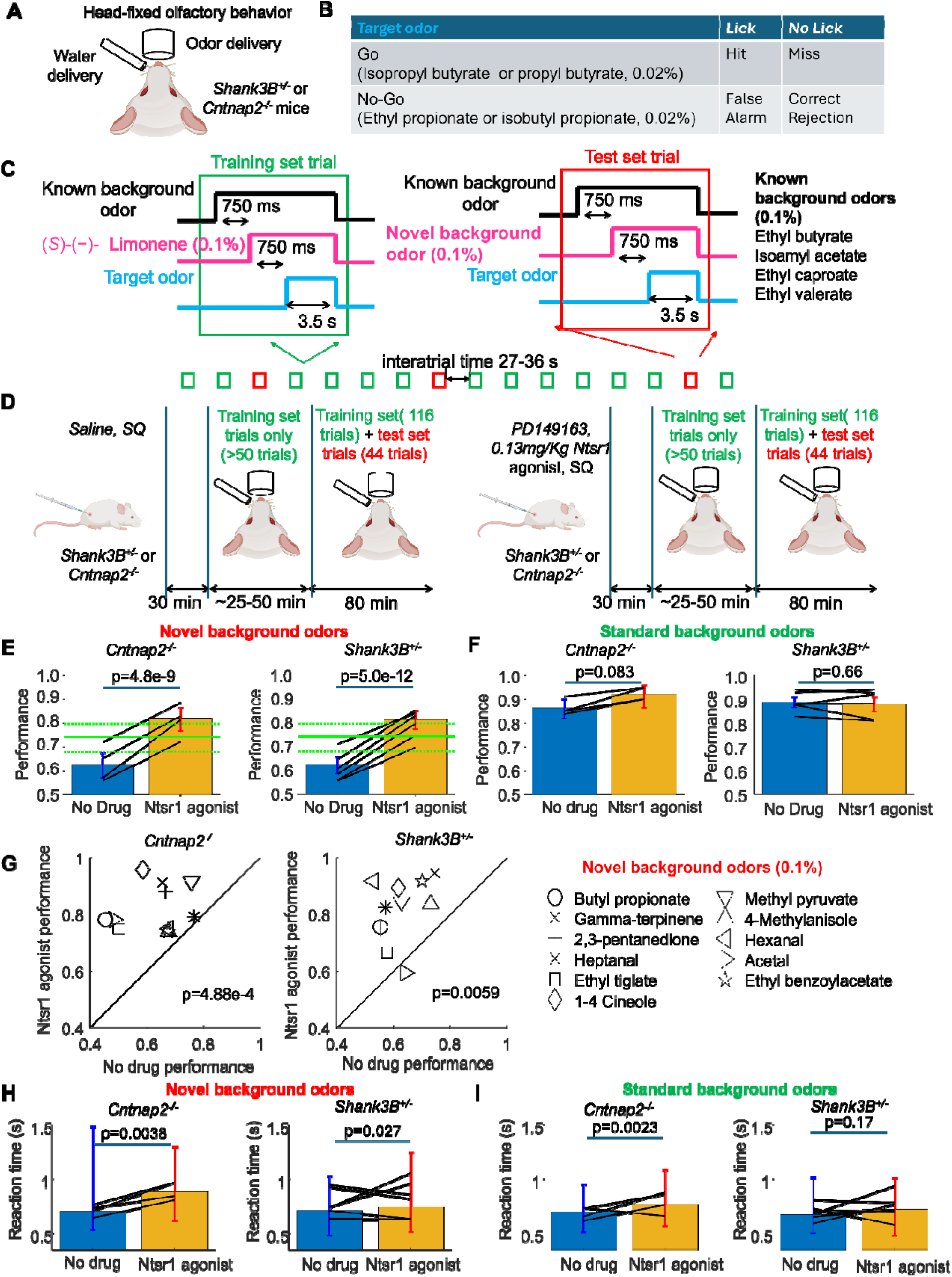
Systemic activation of Neurotensin Receptor 1 rescued target recognition in novel background odors in *Cntnap2*^−/−^ and *Shank3B*^+/−^ mice. (A,B) Head-fixed water-deprived mice were trained in a go/no-go odor recognition task. Mice licked in response to either of two go target odors to receive a water reward and withheld licking in response to either of two no-go target odors.(C) During training, target odors were presented in the presence of familiar background odor mixtures consisting of (S)-(−)-limonene and one of four variable background odors. During testing, (S)-(−)-limonene was replaced on some trials by one of 11 novel background odors that had not been encountered during training. (D) Experimental design. Mice received subcutaneous injections of saline or the Ntsr1 agonist PD149163 30 min before testing. (E) Performance on novel-background trials. Subcutaneous administration of the Ntsr1 agonist improved performance in both *Cntnap2*^−/−^ (n = 4 mice) and *Shank3B*^+/−^ (n = 6 mice) mice. Black lines connect the mean performance of individual mice under saline and drug conditions. Bars indicate group mean ± 95% confidence interval. Green symbols indicate performance of wild-type mice for reference. (F) Performance on familiar-background trials. Ntsr1 agonist administration did not significantly alter performance for familiar background odors.(G) Performance for individual novel background odors. Improvement following Ntsr1 activation was distributed across most novel odors in both genotypes. (H) Reaction times on correct go trials (hits) during novel-background conditions. Ntsr1 agonist administration increased reaction times in both mouse models. Bars indicate median reaction time and error bars denote the 10th–90th percentile range. Black lines connect individual mice. (I) Reaction times during familiar-background trials. Effects of Ntsr1 agonist administration on reaction time were smaller and genotype dependent.

Briefly, water-deprived *Cntnap2*^−/−^ and *Shank3B*^+/−^ mice of both sexes (see **Table 1** for list of animals used) were trained to identify weak target odors in the presence of strong background odors ^5^. Head-fixed mice were trained to lick in response to go target odors (selected from one of two possible odors presented at 0.02% vapor pressure) to receive a 4 μL water reward and to withhold licking in response to no-go target odors (selected from one of two possible odors presented at 0.02% vapor pressure) to avoid a timeout.

Target odors were presented together with two background odors. During the training phase, one background odor was randomly selected from a set of four odors and presented at 0.1% vapor pressure beginning 1.5 s before target odor onset (known background odors). A second background odor, (S)-(-)-limonene, was presented at the same concentration beginning 0.75 s before target onset. During training-set trials, only 8 of the 16 possible combinations of target odors and known background odors were presented. For the testing-set trials, the target odors and the four known background odors were unchanged. Novelty was introduced by replacing (S)-(-)-limonene with one of 11 odors that had not been encountered during training. Testing-set trials also included the 8 target/background combinations that were not presented during training.

Each testing session began with training-set trials containing only the trained background odor combinations with (S)-(-)-limonene. This initial block ensured that animals were motivated and performed the learned odor discrimination task at a high level before novel background odors were introduced. Thus, poor performance on novel-background trials could not be attributed to a failure to perform the task or a lack of motivation on that testing day. Once performance exceeded 80% correct over at least 50 trials, novel-background trials were randomly interleaved with training-set trials. Novel-background trials represented 44 of 160 trials (27.5%) in each session. Each of the 11 novel odors was presented no more than four times per day. On each novel-background trial, the target odor and known background odor were selected randomly, and successive presentations of the same novel odor were separated by at least 30 min. This sparse and variable presentation schedule minimizes learning of the novel background odors over time^6^. By using 11 different novel background odors delivered with an automated odor machine^5^, this task provides a robust and quantitative measure of odor recognition under conditions of novelty. The large number of novel odors allows novelty responses to be characterized reliably within individual mice while minimizing the contribution of odor-specific effects. Automated odor delivery also provides precise control over odor concentration and timing. In contrast, traditional habituation/dishabituation assays that have been used in mouse models of ASD^43,44^ are manually administered and typically measure responses to only a few odors per animal, limiting their ability to distinguish general novelty-processing deficits from odor-specific effects.

*Cntnap2*^−/−^ (n = 4) and *Shank3B*^+/−^ (n = 6) mice received subcutaneous injections of the Ntsr1 agonist PD149163 dissolved in 0.9% saline (0.9% NaCl; 40 μg/mL, 42.4 μM) at a dose of 0.13 mg/kg 30 min before behavioral experiments (see **Figure 1D**). Each mouse was tested on two Ntsr1 agonist treatment days, separated by a saline-control day in which the same injection volume of vehicle was administered. In addition, mice were tested on 2–3 sessions containing novel background odors without any injections.

Systemic administration of the Ntsr1 agonist markedly improved recognition of target odors presented in novel background odors in both ASD mouse models (see **Figure 1E**). The average performance of *Cntnap2*^−/−^ mice without the Ntsr1 agonist was 62.3% (334 trials, 15 sessions, 4 mice). Following administration of the Ntsr1 agonist, performance significantly improved (p=4.8×10⁻, Fisher exact test) to 81.9% (260 trials, 7 sessions). The increase in performance for individual animals (18.8% ± 3.7%, mean ± s.d.) was positive in all 4 *Cntnap2*^−/−^ mice and was significant (p=0.0021, double-tailed paired t-test).

Similarly, the average performance of *Shank3B*^+/−^ mice without the Ntsr1 agonist was 62.2% (754 trials, 24 sessions). Performance also significantly improved (p=5.0×10⁻¹², Fisher test) to 81.3% (418 trials, 12 sessions, 6 mice) following Ntsr1 agonist administration. The increase in performance for individual animals (18.3% ± 3.9%) was positive in all 6 tested *Shank3B*^+/−^ mice and was significant (p=8.7×10⁻, double-tailed paired t-test).

Performance on trials with known background odors was not reduced by the Ntsr1 agonist (see **Figure 1F**). For *Shank3B*^+/−^ mice, performance was 88.6% (863 trials) without the Ntsr1 agonist and remained similar at 87.8% (500 trials) following Ntsr1 agonist administration. This difference was not significant (p=0.66, Fisher exact test). For *Cntnap2*^−/−^ mice, performance was 86.3% (300 trials) without the Ntsr1 agonist and increased to 92.1% (140 trials) following Ntsr1 agonist administration; however, this increase was not significant (p=0.083, Fisher exact test). Thus, *Cntnap2*^−/−^ and *Shank3B*^+/−^ mice maintained high performance on trials with known background odors during Ntsr1 agonist treatment, while showing improved performance on trials with novel background odors.

We next asked whether the improvement in performance produced by the Ntsr1 agonist was broadly distributed across novel background odors or restricted to a small subset of odors (see **Figure 1G**). Because each novel background odor was presented only a few times in each animal (<9 total presentations), data were pooled across animals for this analysis. Performance increased for all 11 novel background odors in *Cntnap2*^−/−^ mice (p=4.88 × 10⁻, binomial test) and for 10 of 11 novel background odors in *Shank3B*^+/−^ mice (p=0.0059, binomial test). Thus, the improvement associated with Ntsr1 agonist administration was distributed across nearly all novel background odors rather than being driven by a small number of novel background odors.

We next asked whether mice learned to recognize the novel background odors during Ntsr1 agonist treatment or whether the improved performance reflected enhanced generalization in the presence of novel background odors. If learning were responsible for the improvement, performance should remain elevated after the drug was discontinued. To test this possibility, we compared performance on the first day of Ntsr1 agonist administration with performance on the following day, when mice instead received saline injections. In *Shank3B*^+/−^ mice, performance on the first day of Ntsr1 agonist administration was 82.1% (207 trials, 6 sessions). On the following day, after saline injection, performance significantly decreased to 71.8% (209 trials, 6 sessions; p = 0.0144, Fisher exact test). Similarly, in *Cntnap2*^−/−^ mice, performance on the first day of Ntsr1 agonist administration was 78.4% (148 trials, 4 sessions). After saline injection the following day, performance significantly decreased to 63.2% (106 trials, 4 sessions; p = 0.011, Fisher exact test). The rapid drop in performance following discontinuation of the Ntsr1 agonist indicates that the improvement cannot be readily explained by learning of the novel background odors but instead reflects enhanced generalization to novel sensory environments.

### Subcutaneous injection of Ntsr1 agonist slowed down reaction time in mouse models of ASD

Generalization to a stimulus that differs from previously learned examples is often associated with longer reaction times while maintaining correct performance^45^, suggesting that additional processing may be required to correctly identify behaviorally relevant stimuli under novel conditions. Similarly, mice performing difficult olfactory discriminations exhibit longer response latencies, and slower reaction times have been associated with improved accuracy^46^. We therefore asked whether the improvement in odor recognition under novel background conditions produced by Ntsr1 activation would also be accompanied by increases in reaction time. To address this question, we analyzed response latencies on hit trials, measured from the onset of the target go odor.

For both *Cntnap2*^−/−^ and *Shank3B*^+/−^ mice, activation of Ntsr1 in the entorhinal cortex significantly increased reaction times during trials with novel background odors (see **Figure 1H**). For *Cntnap2*^−/−^ mice, the median latency from target odor onset was 699 ms (71 trials) [10th–90th percentile: 528–1486 ms] without drug treatment. Ntsr1 agonist administration significantly increased median reaction time (p=0.0038, Wilcoxon rank sum test) to 884 ms (90 trials) [10th– 90th percentile: 610–1298 ms]. For *Shank3B*^+/−^ mice, the median latency was 706 ms (191 trials) [10th–90th percentile: 476–1028 ms] without drug treatment. Ntsr1 agonist administration also significantly increased median reaction time (p=0.027, Wilcoxon rank sum test) to 743 ms (176 trials) [10th–90th percentile: 509–1246 ms]. Interestingly, although the Ntsr1 agonist slowed reaction times, it increased correct lick responses to go target odors presented with novel background odors, arguing against the possibility that the improvement in performance resulted from a general suppression of licking behavior (see **Supplementary Figure 1** and **Supplementary Material: Systemic Ntsr1 activation increased performance on go trials**).

We next examined whether the slowing of reaction times generalized to standard background odors (see **Figure 1I**). In *Cntnap2*^−/−^ mice, median latency was 694 ms (135 trials) [10th–90th percentile: 505–947 ms] without drug treatment and increased significantly to 763 ms (68 trials) [10th–90th percentile: 561–1082 ms] following Ntsr1 agonist administration (p=0.0023, Wilcoxon rank sum test). In *Shank3B*^+/−^ mice, median latency was 669 ms (396 trials) [10th–90th percentile: 495–1011 ms] without drug treatment and was not significantly affected by Ntsr1 agonist administration (717 ms, 223 trials) [10th–90th percentile: 471–1010 ms; p=0.17, Wilcoxon rank sum test].

Overall, Ntsr1 agonist administration increased reaction times for novel background odor trials in both genotypes, while effects on standard odor trials were genotype dependent. Despite the increase in performance produced by subcutaneous injections of the Ntsr1 agonist, there was no increase in basal sniff rate nor in sniff rate during target odor presentation, both of which are elevated when mice are more engaged in a task^47–49^ (see **Supplementary Figure 2** and **Supplementary Material: Systemic Ntsr1 activation did not increase sniff rate in Shank3B+/− and *Cntnap2*^−/−^ mice**). Subcutaneous Ntsr1 agonist administration slightly reduced basal sniff rate in *Cntnap2*^−/−^ mice and did not significantly change sniff responses in *Shank3B*^+/−^ mice.

### Systemic Ntsr1 activation preferentially recruited the entorhinal cortex among Ntsr1-expressing regions

Subcutaneous administration of the Ntsr1 agonist PD149163 improved odor recognition in the presence of novel background odors in both *Cntnap2*^−/−^ and *Shank3B^+/−^* mice. Because PD149163 crosses the blood-brain barrier and reaches the entire brain, the neural substrate responsible for this behavioral rescue is not immediately apparent. To identify candidate regions mediating the behavioral rescue, we first examined the distribution of Ntsr1 expression. Because PD149163 binds selectively to Ntsr1 and shows negligible affinity for other neurotensin receptors or unrelated receptors, including dopaminergic and serotonergic receptors^39^, any brain regions directly driven by PD149163 should express Ntsr1.

Surprisingly, Ntsr1 mRNA expression was low in the main olfactory bulb (MOB) and in several major olfactory bulb target regions, including the piriform cortex (PC), anterior olfactory nucleus (AON), taenia tecta (TT), nucleus of the lateral olfactory tract (NLOT), olfactory tubercle (OT), and cortical amygdala (COA) as determined by in situ hybridization data from the Allen Brain Atlas (see **Figure 2A-C**). Expression was quantified using the Allen Brain Atlas expression energy metric, a measure that incorporates both the density and intensity of Ntsr1 mRNA labeling within each brain region (see Methods). Thus, despite the improvement in odor recognition, the primary olfactory pathway is unlikely to be the direct target of the Ntsr1 agonist (see **Figure 2D**).

**Figure 2.**
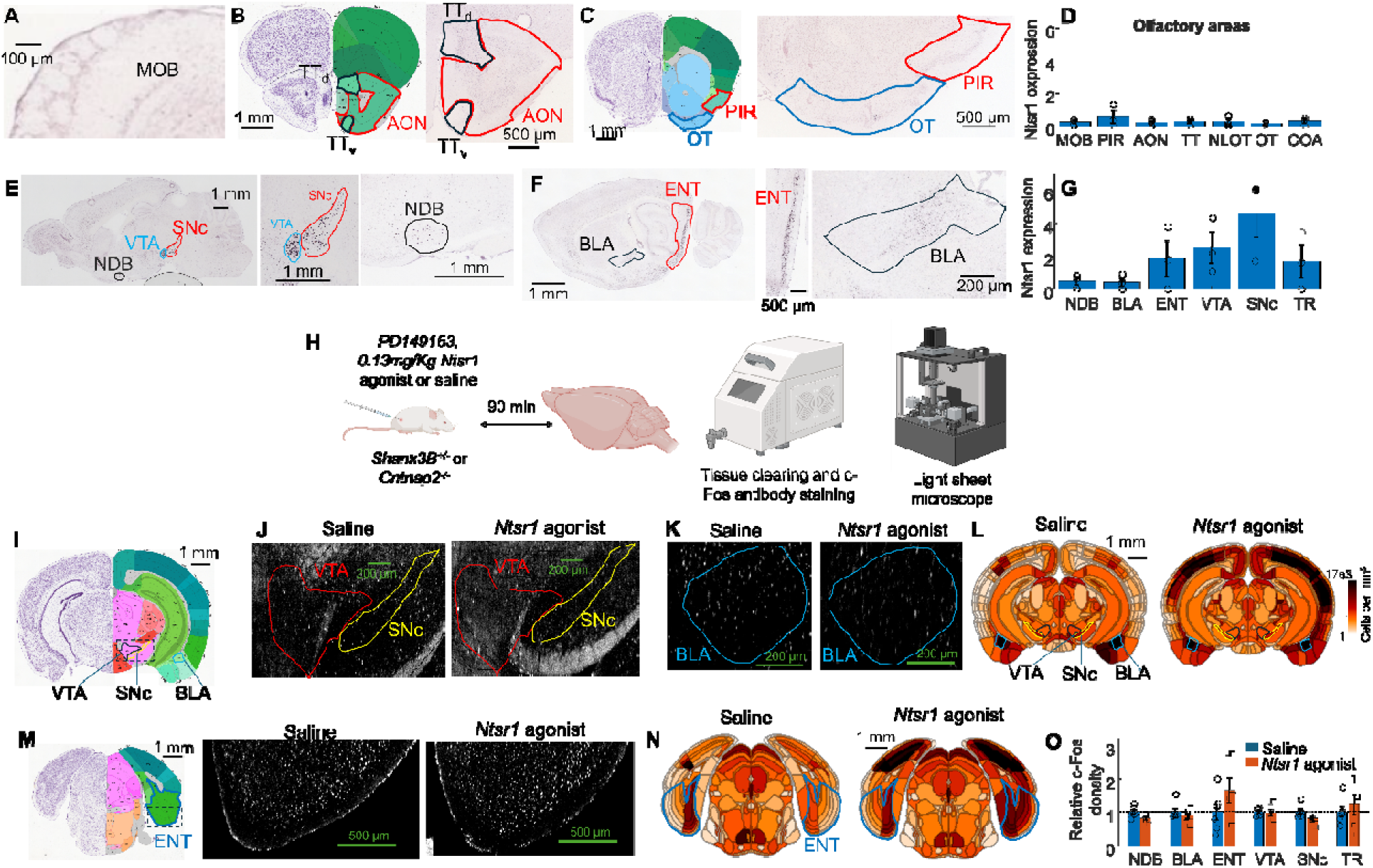
Systemic Ntsr1 activation preferentially recruited the entorhinal cortex among Ntsr1-expressing regions. (A–C) Ntsr1 mRNA expression in olfactory regions using in situ hybridization data from the Allen Brain Atlas. Representative sections of the main olfactory bulb (MOB), anterior olfactory nucleus (AON), taenia tecta (TTd, TTv), piriform cortex (PIR), and olfactory tubercle (OT) show low Ntsr1 expression. (D) Quantification of Ntsr1 expression in regions receiving MOB input (mean ± s.e.m.; n = 3 mice). Symbols represent individual mice. (E–G) Ntsr1 expression is higher in non-olfactory regions. Representative sections show high Ntsr1 expression in the entorhinal cortex (ENT), ventral tegmental area (VTA), substantia nigra pars compacta (SNc), nucleus of the diagonal band (NDB), basolateral amygdala (BLA), and post-piriform transition area (TR). (G) Quantification of Ntsr1 expression in selected brain regions (mean ± s.e.m.; n = 3 mice). Symbols represent individual mice. (H) Experimental design. *Cntnap2*^−/−^ (n = 3) and *Shank3B*^+/−^ (n = 3) mice received subcutaneous injections of saline or the Ntsr1 agonist PD149163. Brains were collected 90 min after injection, cleared, and imaged using light-sheet microscopy for whole-brain c-Fos quantification. (I–L) Regions expressing Ntsr1 but showing no increase in c-Fos expression following systemic Ntsr1 agonist administration. Representative sections from *Shank3B*+/− mice (J,K) and group population density maps (L) show no significant increase in c-Fos-positive cells in the BLA, VTA, or SNc. (M–O) Entorhinal cortex activation following systemic Ntsr1 agonist administration. Representative sections (M) and group population density maps (N) show increased c-Fos expression in the entorhinal cortex. (O) Among the Ntsr1-expressing regions, c-Fos induction was observed only in the entorhinal cortex (mean ± s.e.m.; n = 6 hemispheres per condition). Symbols represent individual hemispheres.

Stronger Ntsr1 expression was restricted to a relatively small number of brain regions (see **Figure 2E–G)**. In adult mice (P56 male C57BL/6), the highest expression was observed in the substantia nigra pars compacta (SNc; expression energy = 4.59 ± 1.46, mean ± s.e.m.; n = 3 mice), ventral tegmental area (VTA; 2.52 ± 0.97), entorhinal cortex (ENT; 1.84 ± 1.09), and post-piriform transition area (TR; 1.68 ± 1.00), with lower expression in the nucleus of the diagonal band (NDB) and basolateral amygdala (BLA). All of these regions exhibited expression energies at least threefold higher than the olfactory region with the highest Ntsr1 expression, the piriform cortex (PIR; 0.53 ± 0.39, mean ± s.e.m.; n = 3 mice). These Ntsr1-expressing regions therefore represented plausible sites through which systemic PD149163 could influence odor recognition behavior.

To determine which of these Ntsr1-expressing regions were activated by systemic PD149163 administration, we quantified c-Fos expression using whole-brain tissue clearing and automated cell counting (see **Figure 2H**). Brain-wide c-Fos mapping has previously been used to identify the neural circuits engaged by pharmacological manipulations and to localize candidate sites of drug action^50^. This approach provided an unbiased and quantitative measure of brain-wide activation following PD149163 administration. PD149163 was administered at a dose of 0.13 mg/kg to 3 mice (2 *Shank3B*^+/−^ mice and 1 *Cntnap2*^−/−^ mouse), and c-Fos-positive cell counts were compared with those from 3 saline-injected control mice (2 *Shank3B*^+/−^ mice and 1 *Cntnap2*^−/−^ mouse). Following injection, mice were returned to their home cages for 2 h before perfusion with saline followed by 4% PFA.

For each brain region, c-Fos density was normalized to the average density measured in saline-treated controls. Among the Ntsr1-expressing regions examined, the entorhinal cortex showed the largest increase in c-Fos expression following PD149163 administration (1.64 ± 0.38-fold over saline controls, mean ± s.e.m.; n = 6 hemispheres), whereas c-Fos induction in all other Ntsr1-expressing regions was modest (<1.2-fold over saline controls; see **Figure 2I–O**). Ntsr1 expression was observed in both the medial and lateral entorhinal cortex and increased c-Fos expression was detected in both subdivisions following PD149163 administration (see **Supplementary Figure 3** and see Supplementary Material: **Ntsr1 is expressed in both medial and lateral entorhinal cortex, and systemic Ntsr1 activation increased c-Fos expression in both subdivisions**).

The combination of strong Ntsr1 expression and robust c-Fos activation identified the entorhinal cortex as the most likely mediator of the behavioral effects of systemic PD149163. We therefore hypothesized that direct activation of Ntsr1-expressing neurons in the entorhinal cortex would be sufficient to rescue odor recognition in novel environments.

### Direct activation of Ntsr1-expressing neurons in the entorhinal cortex rescued olfactory behavior in *Shank3B^+/−^* and *Cntnap2^−/−^*mice

To directly test whether activation of Ntsr1-expressing neurons in the entorhinal cortex is sufficient to rescue odor recognition in novel environments, PD149163 was delivered bilaterally into the entorhinal cortex at a concentration of 4.2 nM (1 μL per site over 5 min), approximately 10,000-fold lower than that used for systemic subcutaneous administration. *Shank3B*^+/−^ and *Cntnap2*^−/−^ mice were implanted bilaterally with guide cannulas targeting the entorhinal cortex (see **Figure 3A**). Because Ntsr1 expression is high in the entorhinal cortex and comparatively low in surrounding regions (see **Figure 3B-C**), and PD149163 is a highly selective Ntsr1 agonist^39^, local infusion is expected to preferentially activate Ntsr1-expressing neurons within the entorhinal cortex.

**Figure 3.**
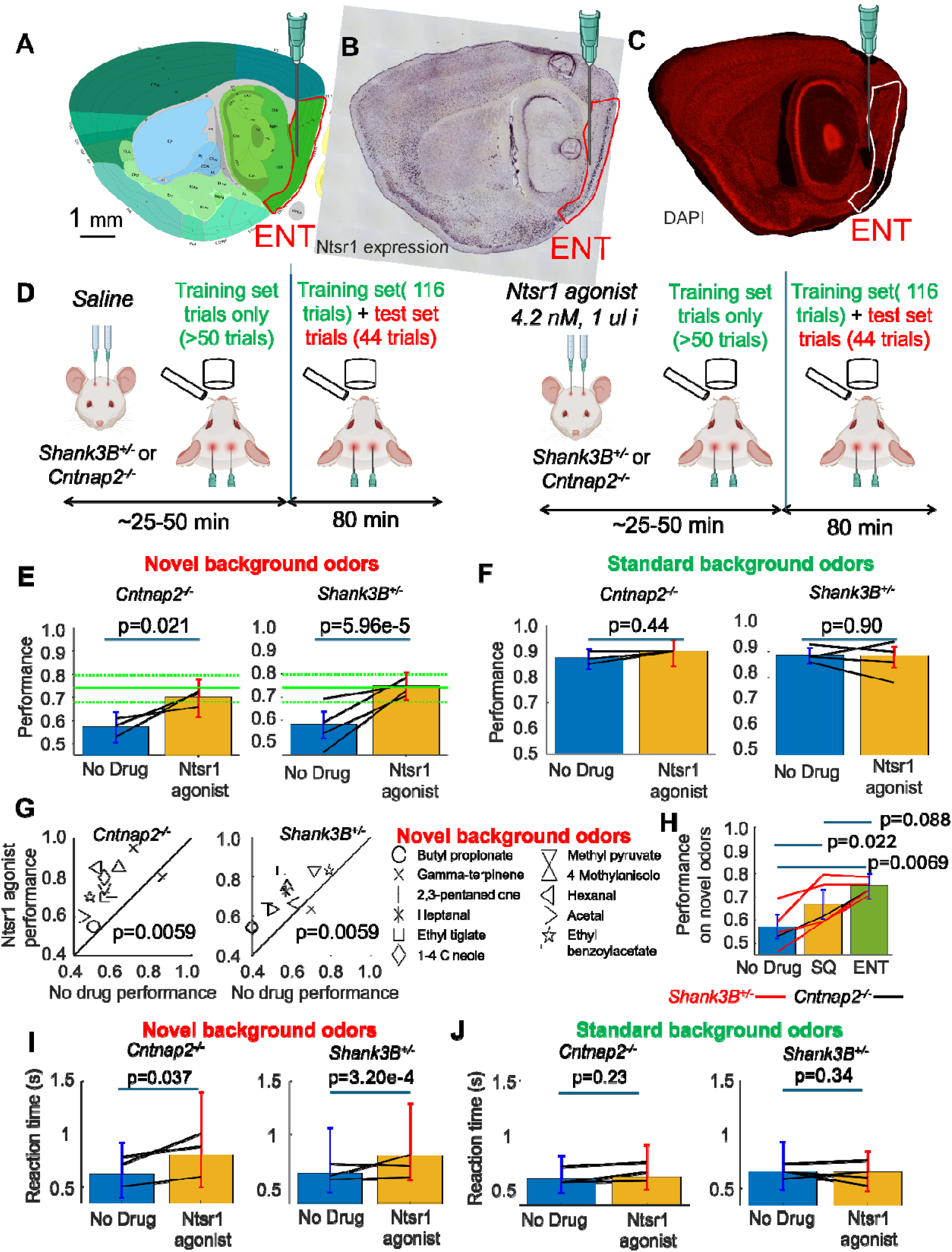
**Direct activation of Ntsr1-expressing neurons in the entorhinal cortex rescued olfactory behavior in *Shank3B*+/− and *Cntnap2*^−/−^ mice**. (A) Cannulas were implanted bilaterally into the entorhinal cortex of *Shank3B*^+/−^ and *Cntnap2*^−/−^ mice. (B) In situ hybridization image showing that Ntsr1 expression is highly enriched in the entorhinal cortex. (C) DAPI image from a *Shank3B*^+/−^ mouse showing the cannula track terminating in the entorhinal cortex. (D) Experimental design. Mice received bilateral entorhinal injections of saline or the Ntsr1 agonist PD149163 before behavioral testing. (E) Performance on novel-background trials following systemic administration. PD149163 significantly improved performance in both *Cntnap2*^−/−^ (n = 3 mice) and *Shank3B*^+/−^ (n = 4 mice) mice. Black lines connect the mean performance of individual mice under saline and drug conditions. Bars indicate the group mean ± 95% confidence interval. Green symbols indicate the performance of wild-type mice for reference.(F) Performance on familiar-background trials. PD149163 did not significantly alter performance on trials containing familiar background odors. (G) Performance for individual novel background odors. Improvement following Ntsr1 activation was observed across most novel background odors in both genotypes. (H) Improvement in performance on novel-background trials was comparable following systemic administration and direct entorhinal infusion of PD149163. (I) Reaction times on correct go trials (hits) during novel-background trials. PD149163 increased reaction times in both mouse models. Bars indicate the median reaction time, and error bars denote the 10th– 90th percentile range. Black lines connect individual mice. (J) Reaction times during familiar-background trials. PD149163 had no significant effect on reaction time.

After training on the odor recognition task, mice (4 *Shank3B*^+/−^ and 3 *Cntnap2*^−/−^ mice) received intra-entorhinal infusions of the Ntsr1 agonist (see **Figure 3D**). Behavioral performance was measured on at least three conditions: a baseline day prior to drug infusion, a drug day, and a post-infusion washout period (≥6 days). A prolonged washout period was required because behavioral effects persisted into the following day and returned to baseline only after ∼6 days (see **Supplementary Figure 4 and** Supplementary Material: **Long lasting behavioral improvement in *Shank3B^+/−^* and *Cntnap2*^−/−^ mice after injection of Ntsr1 agonist in the entorhinal cortex**).

Direct activation of Ntsr1-expressing neurons in the entorhinal cortex was sufficient to improve recognition of target odors in novel sensory environments (see **Figure 3E**). In *Cntnap2*^−/−^ mice, performance without drug was 57.2% (n=222 trials, 6 sessions, 3 mice), and intra-entorhinal PD149163 significantly improved performance to 70.1% (n=127 trials, 3 sessions, 3 mice; p=0.021, Fisher exact test). The increase was observed in all 3 mice (12.9% ± 7.5%, mean ± s.d.) and was significant at the animal level (p=0.048, single-tailed paired t-test).

In *Shank3B*^+/−^ mice, performance on novel background odor trials without drug was 58.1% (n=298 trials, 8 sessions, 4 mice). Intra-entorhinal PD149163 significantly improved performance to 75.0% (n=220 trials, 5 sessions, 4 mice; p=5.96×10⁻, Fisher exact test). The increase was observed in all 4 mice (17.0% ± 8.7%) and was significant at the animal level (p=0.015, single-tailed paired t-test).

Importantly, performance on trials with familiar background odors was unaffected by intra-entorhinal PD149163, consistent with the effects observed during systemic drug administration (see **Figure 3F**). In *Shank3B*^+/−^ mice, performance was 88.8% (400 trials) without drug and 88.4% (250 trials) with drug (p=0.90, Fisher exact test). In *Cntnap2*^−/−^ mice, performance was 87.3% (300 trials) without drug and 90.0% (150 trials) with drug (p=0.44, Fisher exact test). Thus, entorhinal Ntsr1 activation selectively improved performance under novel conditions without compromising performance in familiar conditions, indicating that the rescue was not achieved at the expense of previously learned odor discriminations.

The improvement generalized across most novel background odors (see **Figure 3G**). Performance increased for 10/11 odors in *Cntnap2*^−/−^ mice (p=0.0059, binomial test) and 10/11 odors in *Shank3B*^+/−^ mice (p=0.0059, binomial test), indicating that the effect was broadly distributed across odor identity rather than driven by a subset of stimuli.

To assess whether the observed effects could be explained by learning of novel background odors across days, we compared performance on drug-free sessions before and after entorhinal infusion. In *Cntnap2*^−/−^ mice, performance was 59.3% (n=108 trials, 3 sessions, 3 mice) before drug infusion and 55.3% (n=114 trials, 3 sessions) after 6.3 ± 1.2 days, with no significant difference (p=0.59, Fisher exact test). In *Shank3B*^+/−^ mice, performance was 60.1% (n=143 trials, 4 sessions, 4 mice) before drug infusion and 56.1% (n=155 trials, 4 sessions) after 7.0 ± 2.8 days, also with no significant difference (p=0.56, Fisher exact test). Thus, behavioral improvements following intra-entorhinal PD149163 cannot be explained by learning across days.

### Entorhinal Ntsr1 activation recapitulated the behavioral rescue produced by systemic PD149163

Direct activation of Ntsr1-expressing neurons in the entorhinal cortex rescued odor recognition in novel background odors, similar to the behavioral rescue produced by systemic administration of Ntsr1 agonist. However, Ntsr1 is also expressed in several other brain regions, including the ventral tegmental area (VTA), substantia nigra pars compacta (SNc), post-piriform transition area (TR), nucleus of the diagonal band (NDB), and basolateral amygdala (BLA). Although these regions showed low c-Fos activation following systemic Ntsr1 agonist administration (see **Figure 2D**), it remained possible that they contributed to the behavioral rescue produced by systemic Ntsr1 activation.

If activation of these additional Ntsr1-expressing regions contributes substantially to the rescue, systemic administration of PD149163 would be expected to produce a larger improvement in performance than direct infusion into the entorhinal cortex alone. To test this possibility, a subgroup of 5 mice (4 *Shank3B*^+/−^ and 1 *Cntnap2*^−/−^ mouse) implanted with entorhinal cannulas also received systemic PD149163 (0.13 mg/kg, subcutaneous) on a separate testing day (see **Figure 3H**).

For this subgroup, performance on novel background odor trials in the absence of Ntsr1 agonist was 57.1% (n=368 trials, 10 sessions, 5 mice). Systemic administration improved performance to 66.8% (n=220 trials, 5 sessions). The improvement was positive in all 5 mice (10.5% ± 6.4%, mean ± s.d.) and was significantly greater than zero (p=0.022, paired t-test).

Direct infusion of Ntsr1 agonist into the entorhinal cortex produced a performance of 74.6% (n=264 trials, 6 sessions). The improvement was also positive in all 5 mice (17.5% ± 7.7%, mean ± s.d.) and was significantly greater than zero (p=0.0069, paired t-test).

Importantly, performance following direct entorhinal infusion was not significantly different from performance following systemic administration (p=0.088, paired t-test, n=5 mice). Thus, recruitment of additional Ntsr1-expressing brain regions by systemic PD149163 did not produce a measurable improvement beyond that achieved by activation of the entorhinal cortex alone. These results identify the entorhinal cortex as the principal locus through which Ntsr1 activation improves odor recognition in novel background odors.

### Activation of Ntsr1 in the entorhinal cortex slowed down reaction times for novel background odors

Systemic delivery of the Ntsr1 agonist slowed reaction times for novel background odors in both *Shank3B*^+/−^ and *Cntnap2*^−/−^ mice. Because direct activation of Ntsr1 in the entorhinal cortex reproduced the behavioral rescue produced by systemic PD149163, we next asked whether it would also reproduce the increase in reaction time. If the slowing of reaction times observed after systemic administration is mediated by the entorhinal cortex, then direct activation of entorhinal Ntsr1 should similarly increase response latencies.

Direct activation of Ntsr1 in the entorhinal cortex slowed reaction times for novel background odors in both mouse models (see **Figure 3I**). In *Cntnap2*^−/−^ mice, the median latency from target odor onset was 609 ms (48 trials) [10th–90th percentile: 387–915 ms] without drug treatment. Entorhinal delivery of the Ntsr1 agonist significantly increased the median lick latency (p=0.037, Wilcoxon rank-sum test) to 798 ms (16 trials) [10th–90th percentile: 484–1394 ms].

In *Shank3B*^+/−^ mice, the median latency from target odor onset was 627 ms (106 trials) [10th– 90th percentile: 448–1056 ms] without drug treatment. Entorhinal delivery of the Ntsr1 agonist also significantly increased the median reaction time (p=3.20 × 10⁻, Wilcoxon rank-sum test) to 797 ms (77 trials) [10th–90th percentile: 562–1289 ms].

Thus, activation of Ntsr1 in the entorhinal cortex reproduced the slowing of reaction times observed following systemic administration of PD149163. Importantly, despite the increase in reaction time, correct lick responses to go target odors presented with novel background odors were not significantly reduced (see **Supplementary Figure 5** and Supplementary Material: **Entorhinal injections of Ntsr1 agonist did not reduce correct responses on go trials in *Shank3B*^+/−^ and *Cntnap2*^−/−^ mice**). These results argue against the possibility that the behavioral rescue resulted from a generalized suppression of licking behavior.

Notably, the increase in reaction time with entorhinal Ntsr1 activation was specific to novel background odors and was not observed for trials containing familiar background odors (see **Figure 3J**). In *Shank3B*^+/−^ mice, the median latency from target odor onset was 653 ms (190 trials) [10th–90th percentile: 478–933 ms] without drug treatment. Entorhinal delivery of the Ntsr1 agonist did not significantly alter the median latency (p=0.34, Wilcoxon rank-sum test), which remained at 646 ms (111 trials) [10th–90th percentile: 469–842 ms]. In *Cntnap2*^−/−^ mice, the median latency from target odor onset was 591 ms (143 trials) [10th–90th percentile: 459– 803 ms] without drug treatment. Entorhinal delivery of the Ntsr1 agonist did not significantly change median reaction time (p=0.23, Wilcoxon rank-sum test), which remained at 606 ms (58 trials) [10th–90th percentile: 486–899 ms].

Therefore, activation of Ntsr1 in the entorhinal cortex selectively increased reaction times during odor recognition in novel environments while leaving responses to familiar odor backgrounds unchanged. This contrasts with systemic Ntsr1 activation, which also slowed reaction times for familiar background odors in a genotype-dependent manner (see **Figure 1I**). The broader effects of systemic administration are likely mediated by activation of Ntsr1-expressing regions outside the entorhinal cortex. In contrast, the selective slowing observed following entorhinal Ntsr1 activation occurred under the same conditions in which behavioral performance improved, suggesting that increased reaction times are specifically linked to successful generalization in novel sensory environments.

Consistent with systemic Ntsr1 activation, entorhinal activation did not increase either basal sniff rate or odor-evoked sniff rate (see **Supplementary Figure 6** and Supplementary Material: **Entorhinal injections of Ntsr1 agonist did not increase sniff rate in *Shank3B*^+/−^ and *Cntnap2*^−/−^ mice**). Thus, the behavioral effects of entorhinal Ntsr1 activation cannot be explained by changes in odor sampling.

### Ntsr1 activation preferentially recruited olfactory feedback pathways rather than hippocampal feedforward pathways

We next asked how activation of Ntsr1-expressing neurons in the entorhinal cortex improved generalization to novel sensory environments. One possibility is that entorhinal cortex recruited the hippocampus, which has been shown to be involved in target recognition in novel environments^51^. The entorhinal cortex sends major feedforward projections to the hippocampus and can modulate hippocampal activity^52,53^.

An alternative possibility is that the entorhinal cortex improved generalization by refining sensory representations within the olfactory system. By reducing the impact of novel background odors on neural activity, entorhinal output could generate target representations that are more resistant to interference from unfamiliar stimuli, thereby facilitating generalization. The entorhinal cortex sends feedback projections to the main olfactory bulb (MOB) ^54,55^ and to areas that are reciprocally connected with the MOB, including the piriform cortex and anterior olfactory nucleus (AON). In addition, entorhinal activation can modulate odor responses within these regions^56–59^.

To quantify the relative anatomical strength of feedforward hippocampal and olfactory feedback pathways, we analyzed anterograde tracing data obtained following AAV-EGFP injections into the entorhinal cortex from the Allen Mouse Brain Connectivity Atlas (see **Figure 4A-B**). The strongest relative projection target was the hippocampus, which received 58.7% ± 5.8% of labeled axons (mean ± s.e.m., n=11 mice) (see **Figure 4C**). In comparison, olfactory areas received fewer labeled axons, with the strongest recipients being the piriform cortex (19.5% ± 2.5%), anterior olfactory nucleus (8.1% ± 1.4%), and olfactory tubercle (8.2% ± 1.4%) (see **Figure 4D-F**). Given the dominance of the hippocampal projection (see **Figure 4G-H**), we asked whether Ntsr1 activation would preferentially recruit the hippocampus rather than olfactory areas.

**Figure 4.**
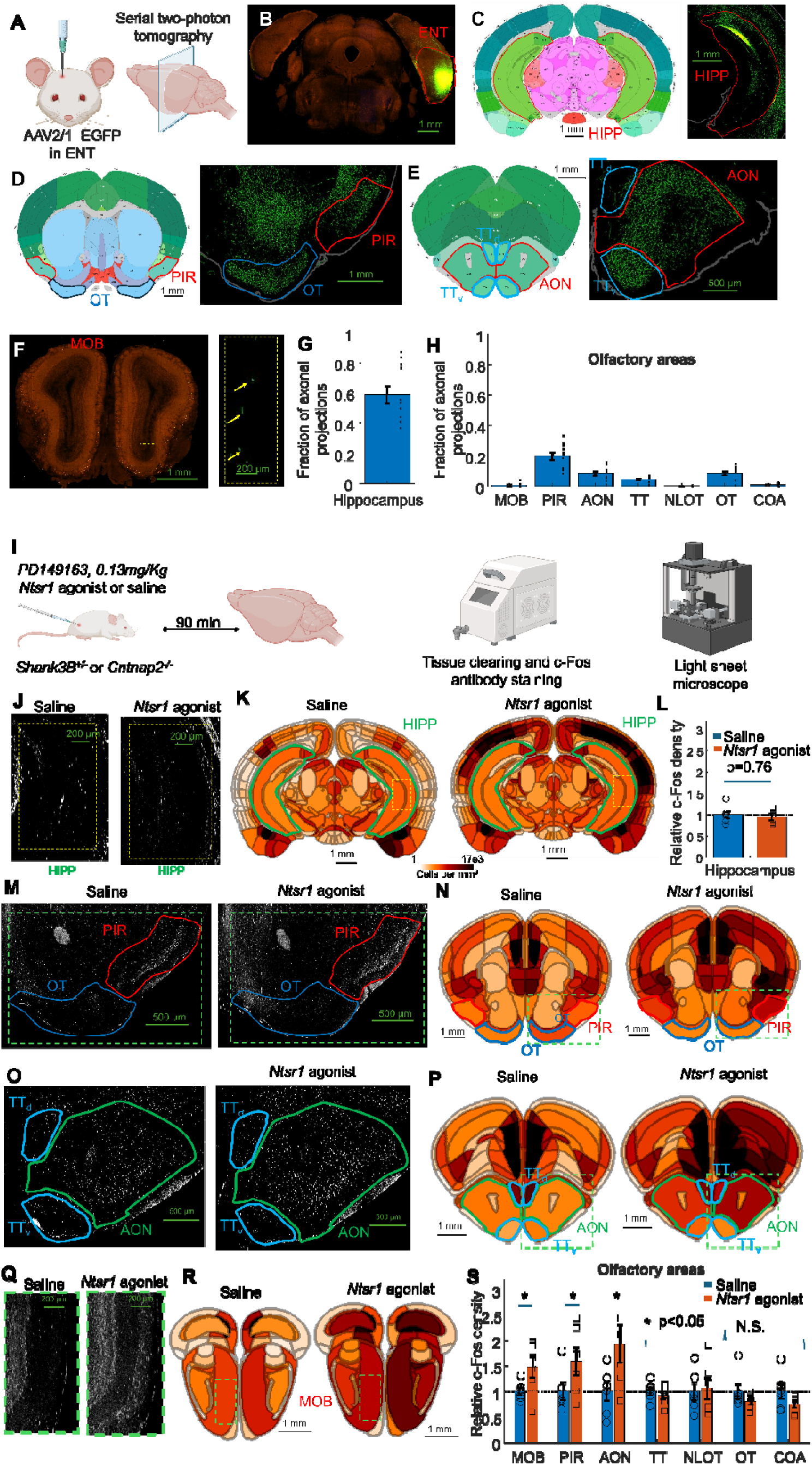
Ntsr1 activation preferentially recruited olfactory feedback pathways rather than hippocamp.al feedforward pathways. ((A–H) Anatomical projections from the entorhinal cortex. (A) Wild-type mice received entorhinal injections of AAV2/1 driving EGFP expression. Axonal projections were imaged using serial two-photon tomography (Allen Mouse Brain Connectivity Atlas). (B) Representative coronal section showing the entorhinal injection site. (C– F) Representative sections showing dense projections to the hippocampus and olfactory cortical areas, including the olfactory tubercle (OT), piriform cortex (PIR), taenia tecta (TT), and anterior olfactory nucleus (AON), and sparse projections to the main olfactory bulb (MOB; yellow arrows). (G,H) Quantification of axonal projections (mean ± s.e.m.; n = 11 mice) showing substantially stronger projections to the hippocampus than to the MOB or regions receiving MOB input. (I–S) Systemic Ntsr1 activation preferentially recruits olfactory feedback pathways. (I) Experimental design. *Cntnap2*^−/−^ (n = 3) and *Shank3B*^+/−^ (n = 3) mice received subcutaneous injections of saline or the Ntsr1 agonist PD149163. Brains were collected 90 min after injection, cleared, and imaged using light-sheet microscopy for whole-brain c-Fos quantification. (J–L) Representative sections (J), average c-Fos density maps (K), and quantification (L) show no increase in hippocampal c-Fos expression following Ntsr1 agonist administration. (M–R) Representative sections and average c-Fos density maps showing increased c-Fos expression in the piriform cortex (PIR), anterior olfactory nucleus (AON), and main olfactory bulb (MOB), but not in the olfactory tubercle (OT) or taenia tecta (TT). (S) Relative c-Fos expression (mean ± s.e.m.; n = 6 hemispheres per condition) showing significant induction in the MOB, PIR, and AON, but not in the hippocampus, OT, or TT. Symbols represent individual hemispheres.

To address this question, we quantified c-Fos expression following subcutaneous administration of the Ntsr1 agonist PD149163 and saline controls (see **Figure 4I**). Despite the dominant projection from entorhinal cortex to hippocampus, hippocampal c-Fos expression (see **Figure 4J-K**) was not significantly increased by Ntsr1 activation (0.94 ± 0.08, p=0.76, single-tailed t-test, n=6) relative to saline controls (see **Figure 4L**). In contrast, significant increases in c-Fos expression were observed in the MOB (1.47 ± 0.21, p=0.038), piriform cortex (1.59 ± 0.28, p=0.044), and AON (1.93 ± 0.38, p=0.029; single-tailed t-tests), indicating selective recruitment of a subset of olfactory circuits (see **Figure 4M-R**), whereas other regions receiving MOB input showed no significant activation (all p>0.35)(see **Figure 4S**). Notably, these increases were observed in untrained mice that were returned to their home cages following drug administration, indicating that recruitment of olfactory circuits did not depend on active odor-guided behavior. This pattern was not explained only by anatomical connectivity. Although the olfactory tubercle received substantially stronger entorhinal projections than the MOB, it showed no increase in c-Fos expression following Ntsr1 activation (0.80 ± 0.06, p=0.99, single-tailed t-test). Unlike the piriform cortex and AON, which both send extensive feedback projections to the olfactory bulb, the olfactory tubercle receives olfactory bulb input but does not project back to the MOB^60^. Thus, Ntsr1 activation preferentially recruited olfactory regions that participate in feedback circuits with the olfactory bulb rather than olfactory regions with the strongest entorhinal input.

Interestingly, increased c-Fos expression was observed in olfactory regions that express little or no Ntsr1 (see **Figure 2D**). In particular, the MOB exhibited elevated c-Fos despite minimal Ntsr1 expression and sparse direct entorhinal input (0.6% ± 0.4% of labeled axons, see **Figure 4F**), suggesting that MOB recruitment occurs indirectly through intermediate olfactory areas such as the piriform cortex and AON. Together with the selective recruitment of the piriform cortex and AON, these findings indicate that systemic Ntsr1 activation engages a distributed olfactory network that includes regions reciprocally connected with the MOB.

### Entorhinal Ntsr1 activation reduced olfactory bulb output in mouse models of ASD

Within the olfactory system, activation was concentrated in regions that participate in feedback interactions with the MOB, including the piriform cortex and AON. Because these feedback pathways could suppress olfactory bulb output ^22–27^, and because successful generalization is associated with reduced glomerular responses to familiarized background odors in *Cntnap2^−/−^* and *Shank3B*^+/−^ mice ^8^, we hypothesized that entorhinal Ntsr1 activation might similarly suppress glomerular responses to novel background odors in mouse models of ASD, thereby reducing interference from distractor stimuli and promoting generalization.

To measure glomerular output, we used Thy1-GCaMP6f mice, which express GCaMP6f in mitral and tufted cells, the principal output neurons of the olfactory bulb (see **Figure 5 A-C**)^61^. These mice were crossed with either *Cntnap2*^−/−^ or *Shank3B*^+/−^ mice as previously described^8^. We recorded from two *Cntnap2*^−/−^ × Thy1-GCaMP6f mice and two *Shank3B*^+/−^ × Thy1-GCaMP6f mice implanted with bilateral cannulas targeting the entorhinal cortex. Glomerular activity was compared between sessions in which PD149163 was infused bilaterally into the entorhinal cortex (4.2 nM, 1 μL per site over 5 min) and sessions in which mice received saline infusions or no infusion. We recorded 5 sessions with Ntsr1 agonist infusion and 15 sessions without infusion (see **Figure 5D**).

**Figure 5.**
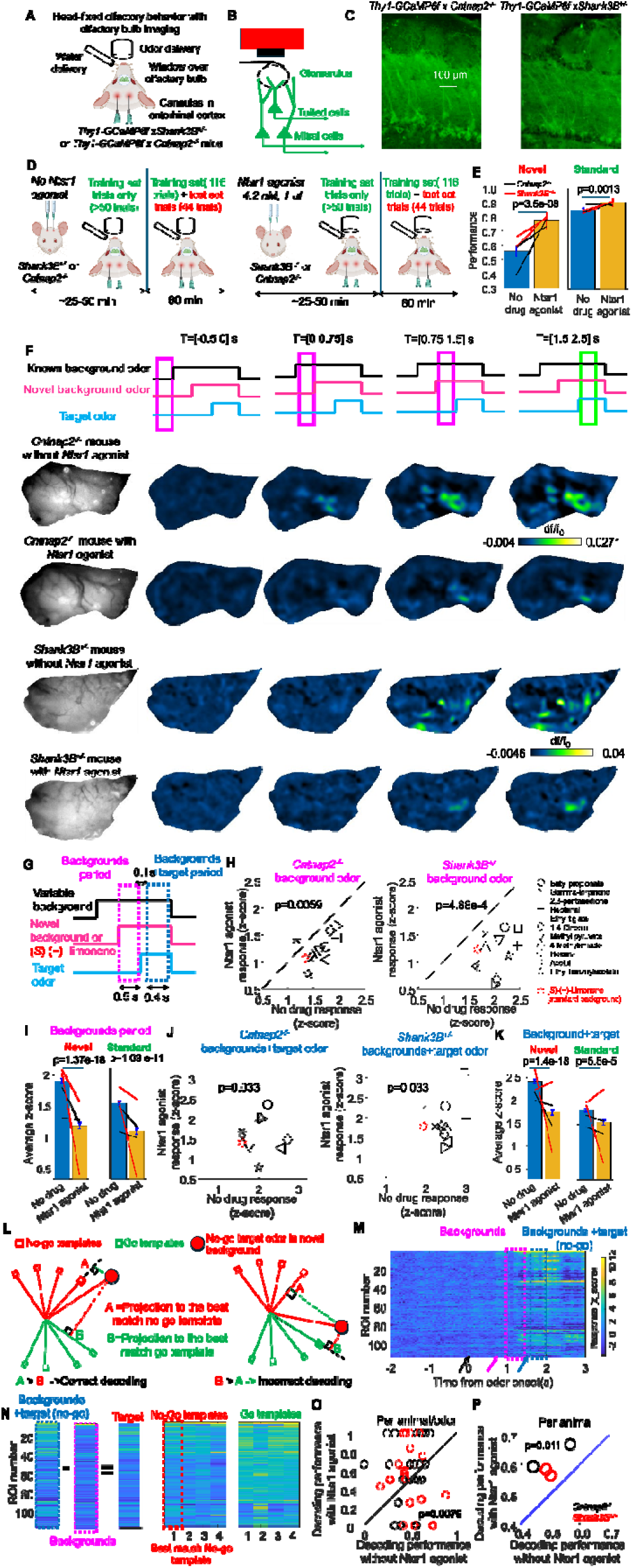
Entorhinal Ntsr1 activation enhanced neural representations that support odor generalization. (A) Schematic of the head-fixed preparation used for widefield calcium imaging during odor discrimination. (B) Schematic of GCaMP6f expression in mitral and tufted cells of Thy1-GCaMP6f mice. (C) Representative histological sections from *Shank3B*^+/−^ and *Cntnap2*^−/−^ mice showing GCaMP6f expression in mitral and tufted cells. (D) Experimental design. *Shank3B*^+/−^ and *Cntnap2*^−/−^ mice implanted with bilateral entorhinal cannulas received no injection or PD149163 before imaging. (E) Entorhinal Ntsr1 activation improved behavioral performance during novel-background trials in imaging sessions. Bars represent the mean, error bars indicate the 95% confidence interval, and black lines connect measurements from individual mice. (F) Representative odor-evoked glomerular activity from single trials showing reduced olfactory bulb responses following entorhinal Ntsr1 activation in a *Cntnap2*^−/−^ mouse and a *Shank3B*^+/−^ mouse. The familiar background odor was ethyl butyrate and the no-go target odor was propyl butyrate. The novel background odor was ethyl benzoylacetate for the *Cntnap2*^−/−^ mouse and acetal for the *Shank3B*^+/−^ mouse. (G) Time windows used for analysis relative to background and target odor presentation. (H) Background-evoked glomerular activity was reduced for most of the 11 novel background odors in both *Shank3B*^+/−^ and *Cntnap2*^−/−^ mice following entorhinal Ntsr1 activation. (I) Mean background-period glomerular activity for novel and familiar background odors in individual animals. Bars represent the mean ± s.e.m. (J) Target-period glomerular activity was reduced for most of the 11 novel background odors following entorhinal Ntsr1 activation. (K) Mean target-period glomerular activity for novel and familiar background odors in individual animals. Bars represent the mean ± s.e.m. (L) Schematic illustrating the neural decoding analysis. Generalization was defined as increased similarity of target odors presented with novel background odors to previously learned odor mixtures associated with the same behavioral valence (go or no-go). Similarity between trial-by-trial responses and the templates was quantified using the dot product. (M) Example glomerular population response during target odor presentation. The black arrow indicates the onset of the familiar variable background odor (ethyl valerate), the magenta arrow indicates the onset of the novel background odor (2,3-pentanedione), and the blue arrow indicates the onset of the no-go target odor (ethyl propionate). (N) Schematic of the neural decoding analysis. Templates were defined as the normalized average glomerular population response for each target–familiar background combination. For novel-background trials, the background-period population response was subtracted from the target-plus-background response to isolate the target-evoked response. (O) Neural generalization performance for each of the 11 novel background odors in individual animals. Entorhinal Ntsr1 activation improved generalization in 30 of 44 animal–odor pairs. (P) Overall neural generalization accuracy increased following entorhinal Ntsr1 activation in all mice. Symbols represent individual mice.

The imaging cohort reproduced the behavioral rescue observed in the larger behavioral experiments (**see Figure 5E**). Performance on novel-background trials increased from 55.5% (95% CI: 51.6–59.3%, 660 trials) without PD149163 to 77.3% (95% CI: 70.6–83.0%, 189 trials) following entorhinal infusion of the Ntsr1 agonist (p=3.50 × 10⁻, Fisher exact test). Improvement was observed in all four mice (21.6 ± 12.3%, mean ± s.d.; p=0.039, paired t-test). A smaller but significant increase was also observed for familiar-background trials, from 85.0% (1895 trials) to 90.4% (530 trials) (p=0.0013, Fisher exact test).

ROIs were drawn over odor-responsive glomeruli (see Methods). We analyzed 732 ROIs from *Cntnap2*^−/−^ mice without drug treatment and 229 ROIs following PD149163 infusion, as well as 759 ROIs from *Shank3B*^+/−^ mice without drug treatment and 264 ROIs following PD149163 infusion. Because behavioral performance differed between drug and no-drug sessions, neural analyses were restricted to correct trials.

Activation of Ntsr1 in the entorhinal cortex produced a robust suppression of glomerular responses to novel background odors (see **Figure 5F** for example images in a *Cntnap2*^−/−^ and a *Shank3B*^−/−^ mouse). We first examined glomerular responses during the 500 ms period preceding target onset, when only the novel and familiar background odors were present (see **Figure 5G**). Average glomerular activity (see **Figure 5H**) was reduced for 10 of 11 novel odors in *Cntnap2*^−/−^ mice (p=0.0059, binomial test) and for all 11 novel odors in *Shank3B*^+/−^ mice (p=4.88 × 10⁻, binomial test). Across all novel-background trials, mean ROI activity decreased from 1.91 ± 0.04 z-score units (mean ± s.e.m.; n=1491 ROIs) without drug treatment to 1.19 ± 0.05 z-score units (n=493 ROIs) following PD149163 infusion. This reduction was observed in all four mice (see **Figure 5I**).

To account for differences between animals and genotypes, we performed a multivariate linear analysis for the average response per ROI (n=1984 ROIs) using animal identity and genotype as covariates. Entorhinal Ntsr1 activation significantly reduced glomerular responses to novel background odors (p=1.4 × 10⁻¹), whereas neither animal identity (p=0.58) nor genotype (p=0.48) had significant effects.

Interestingly, suppression of glomerular activity was not limited to novel background odors. Responses to familiar background odors were also reduced. Average ROI activity decreased from 1.55 ± 0.03 z-score units (n=1491 ROIs) without PD149163 to 1.11 ± 0.05 z-score units (n=493 ROIs) following infusion. Multivariate analysis again revealed a significant effect of Ntsr1 activation (p=1.09 × 10⁻¹¹), with no significant contributions from animal identity (p=0.31) or genotype (p=0.20).

We next examined glomerular responses during the target period, defined as the 400 ms window beginning 100 ms after target odor onset (see **Figure 5J**). Similar to the effects observed during the background period, entorhinal Ntsr1 activation reduced glomerular activity during target presentation, although effects were more variable across animals. Average responses to novel-background trials were reduced for 9 of 11 novel odors in both *Cntnap2*^−/−^ and *Shank3B*^+/−^ mice (p=0.033, binomial test for both genotypes). Across all ROIs, activity decreased from 2.41 ± 0.05 z-score units (n=1491 ROIs) to 1.74 ± 0.07 z-score units (n=493 ROIs). Multivariate analysis revealed a significant effect of Ntsr1 activation (p=5.18 × 10⁻¹), although significant contributions of animal identity (p=0.0043) and genotype (p=0.0027) were also observed (see **Figure 5K**).

Responses to familiar-background trials during the target period were likewise reduced. Mean ROI activity decreased from 1.81 ± 0.04 z-score units (n=1491 ROIs) without drug treatment to 1.54 ± 0.06 z-score units (n=493 ROIs) following PD149163 infusion. Multivariate analysis again identified a significant effect of Ntsr1 activation (p=5.5 × 10⁻), together with significant effects of animal identity (p=0.013) and genotype (p=0.0038).

Thus, activation of Ntsr1 in the entorhinal cortex broadly reduced olfactory bulb output in both *Cntnap2*^−/−^ and *Shank3B*^+/−^ mice. The strongest and most consistent effect occurred during presentation of the background odors, before target onset, where suppression was observed for nearly every novel odor tested and across all animals. These results suggest that entorhinal Ntsr1 activation improves odor recognition in novel environments by reducing glomerular responses evoked by background odors before target odor presentation. Because novel background odors distort neural representations away from those associated with familiar odor combinations, suppressing these responses may make target-containing mixtures more similar to previously learned representations, thereby promoting successful generalization to novel sensory environments.

### Entorhinal Ntsr1 activation enhanced neural representations that support generalization

Activation of Ntsr1-expressing neurons in the entorhinal cortex reduced glomerular output in both *Shank3B*^+/−^ and *Cntnap2*^−/−^ mice for most novel background odors. We next asked whether this suppression altered olfactory bulb representations in a manner that would facilitate generalization of target odor identity to novel sensory environments. Specifically, we tested whether entorhinal Ntsr1 activation transformed glomerular output patterns evoked by novel odor mixtures into representations that more closely resembled those associated with previously learned odor mixtures.

We defined improved generalization as a condition in which the representation of a target odor presented with a novel background becomes more similar to previously learned representations associated with the same behavioral valence (go or no-go) than to representations associated with the opposite valence (see **Figure 5L**). Thus, successful generalization occurs when novel background odors have a reduced impact on the neural representation of the target stimulus, permitting correct decoding of the target’s behavioral valence.

To quantify this, we constructed population vectors from the activity of responsive glomeruli (see **Figure 5M** for an example population response) and measured similarity using the dot product between trial-by-trial responses and odor “templates.” Templates were defined as the average population response for each target–familiar background combination, normalized to unit length. For novel-background trials, the population response during the background period was subtracted from the target+background response to isolate stimulus-evoked activity (see **Figure 5N**).

To determine whether entorhinal Ntsr1 activation improved neural generalization, we quantified how accurately the behavioral valence (go or no-go) of target odors could be decoded on novel-background trials. To account for differences in reaction time induced by Ntsr1 activation and to avoid responses related to licking^8^, decoding was performed using a time window beginning 100 ms after target onset and ending 150 ms before the median lick time. This window was applied to both go and no-go trials and computed separately for standard- and novel-background conditions. A novel-background trial was considered successfully generalized if the best-matching template corresponded to the correct stimulus valence. Only correct behavioral trials were used for template construction and decoding analyses.

For each animal, neural generalization performance was evaluated across the 11 novel background odors (see **Figure 5O**). Across animals and odors, generalization improved for 14/22 animal–odor pairs in *Cntnap2*^−/−^ mice and 16/22 animal–odor pairs in *Shank3B*^+/−^ mice following entorhinal Ntsr1 activation. A mixed-effects linear model including genotype as a covariate revealed no significant effect of genotype (p = 0.37). Overall, Ntsr1 activation improved generalization performance in 68.2% of animal–odor pairs, a rate significantly greater than expected by chance (p = 0.0079, binomial test; 44 total pairs across 4 mice).

Across all mice, generalization accuracy increased significantly following entorhinal Ntsr1 activation, from 49.8%± 5.21% to 60.2% ± 4.5% (see **Figure 5P**)(mean ± s.d.; n = 4 mice; p = 0.011, paired t-test). Thus, activation of Ntsr1 in the entorhinal cortex reduced the influence of novel background odors on glomerular representations and increased the similarity of target-containing mixtures to previously learned representations associated with the same behavioral valence. These changes improved neural generalization and likely contributed to the enhanced recognition of target odors in novel sensory environments.

## Discussion

Successful generalization requires extracting behaviorally relevant information while minimizing interference from unfamiliar background stimuli. ASD is associated with deficits in generalizing learned information to novel contexts^1^, and it has been proposed that unpredictable stimuli may exert an abnormally strong influence on perception because they violate sensory expectations in ASD^2^. Here, we found that activation of Ntsr1-expressing neurons in the entorhinal cortex rescued a deficit in odor recognition under novel sensory conditions in two distinct mouse models of ASD. The rescue was observed following both systemic administration and direct entorhinal infusion of the Ntsr1 agonist PD149163, identifying the entorhinal cortex as a key locus through which neurotensin signaling improves behavioral performance. These findings reveal a previously unrecognized role for entorhinal Ntsr1 signaling in promoting successful generalization to novel sensory environments in mouse models of ASD. The behavioral rescue was accompanied by a reduction in neural responses to novel distractor odors, suggesting that improved generalization can be achieved by reducing the influence of unfamiliar sensory stimuli on neural representations of behaviorally relevant cues.

Although we initially expected Ntsr1 activation to act primarily through the piriform cortex, based on the use of an Ntsr1-Cre line to manipulate piriform corticobulbar feedback^22^ and that the piriform cortex is altered across multiple mouse models of ASD^62^, our anatomical and functional results instead identified the entorhinal cortex as the principal site of action for the Ntsr1 agonist. Despite receiving input from relatively few olfactory bulb neurons compared to other olfactory regions^63^, the entorhinal cortex does exert a disproportionate influence on odor-guided behavior. Inactivation of fan cells in the lateral entorhinal cortex impairs learning of novel odors while sparing discrimination of familiar odors^64^, whereas broader entorhinal inactivation disrupts odor discrimination even for simple odor stimuli ^65^. Beyond olfactory processing, the entorhinal cortex occupies a central position within associative and multimodal networks^66,67^, and entorhinal lesions impair object-context recognition^68^ and spatial navigation^69^. The behavioral rescue observed here may therefore reflect a broader mechanism for improving generalization rather than one specialized for olfactory perception. Because impaired generalization is thought to contribute to behavioral deficits across multiple sensory and cognitive domains in ASD^2^, the effects of entorhinal Ntsr1 activation may have implications beyond olfactory processing given the broad effects of entorhinal cortex activity^53^. Whether entorhinal Ntsr1 activation can facilitate generalization in other sensory modalities in mouse models of ASD remains to be determined.

The entorhinal cortex projects both feedforward to the hippocampus and feedback to olfactory areas, including the piriform cortex and anterior olfactory nucleus (AON)^55^. Surprisingly, Ntsr1 activation preferentially recruited the olfactory areas rather than the hippocampus, despite the hippocampus receiving a larger fraction of entorhinal projections than the olfactory areas. The absence of substantial hippocampal recruitment was unexpected because activation of the entorhinal-hippocampal circuit is strongly associated with memory retrieval and recollection-based performance ^70^, and Ntsr1 activation improved recognition of target odors in novel environments. Instead, improved behavioral performance was accompanied by preferential recruitment of olfactory feedback circuits, including the piriform cortex and AON. These findings suggest that enhancement of odor recognition in novel environments can be achieved through modulation of sensory representations within olfactory circuits without substantial recruitment of hippocampal memory networks.

Both the piriform cortex and the AON provide extensive feedback projections to the olfactory bulb, and activation of these areas is associated with reduced glomerular output^22–27^. Consistent with recruitment of these feedback circuits, entorhinal Ntsr1 activation improved neural generalization in both *Cntnap2*^−/−^ and *Shank3B*^+/−^ mice by suppressing olfactory bulb responses, with the strongest effects occurring during presentation of novel background odors before target onset. This was accompanied by improved decoding of target odor valence and improved behavioral performance in novel sensory environments. By reducing the impact of these novel background stimuli, entorhinal Ntsr1 activation increased the similarity between neural representations of novel and previously learned odor mixtures, thereby restoring successful generalization.

Although corticobulbar feedback suppresses olfactory bulb output, its computational role remains poorly understood. Sparse reconstruction algorithms have been shown to outperform simple linear classifiers at recognizing target odors in the presence of novel background odors ^5^. These algorithms^71–75^ estimate the components of an odor mixture from a large dictionary of possible odor representations and are computationally more demanding than simple pattern classification and can be robust to novel distractors ^76^. One proposed role of cortical feedback is to support such computations by subtracting predictable activity from incoming sensory input and refining sensory representations. Consistent with this framework, entorhinal Ntsr1 activation reduced responses to novel background odors and improved behavioral generalization. Notably, this improvement was accompanied by longer reaction times that were restricted to novel-background trials. Similar increases in reaction time have been observed when humans generalize learned categories to novel stimuli^45^ and when rodents perform more demanding olfactory discriminations^46^. The selective slowing observed here was not accompanied by reductions in licking probability, sniffing behavior, or performance on familiar background odors, arguing against a generalized motor impairment. Instead, the increased reaction times may reflect the additional processing required to correctly interpret novel sensory environments. Consistent with this interpretation, entorhinal Ntsr1 activation simultaneously increased reaction times and improved behavioral performance.

Ntsr1 agonists have been extensively studied for their ability to modulate dopaminergic signaling. Systemic administration of PD149163 reduces amphetamine-induced hyperactivity, a model of schizophrenia^39,40,42^ and Ntsr1 agonists are also being considered as potential treatments for substance use disorder^77^ (Slosky et al 2020). Consistent with these reports, systemic Ntsr1 activation in our experiments produced broader behavioral effects, including increased reaction times during familiar odor discrimination. In contrast, direct infusion of PD149163 into the entorhinal cortex produced a more selective phenotype, improving performance and increasing reaction times specifically during recognition in novel sensory environments. These findings indicate that the enhancement of generalization observed is different from the classical dopaminergic actions of Ntsr1 agonists.

Our results are consistent with previous work showing that Ntsr1 activation in the entorhinal cortex increases neuronal excitability by suppressing TREK-2 potassium channels, producing sustained increases in firing rates^78^ (Xiao et al., 2014). In the same study, entorhinal administration of PD149163 improved spatial learning and memory in an Alzheimer’s disease rodent model. Together with the present findings, these observations suggest that Ntsr1 activation engages entorhinal circuits to enhance performance across multiple cognitive domains, including memory and sensory generalization, and identifies entorhinal Ntsr1 signaling as a potential mechanism for improving generalization in ASD models.

Several limitations should be considered. Although these results identify entorhinal Ntsr1 activation as sufficient to improve generalization in mouse models of ASD, they do not establish which downstream projections are necessary for the behavioral rescue. Disentangling the contributions of piriform cortex and anterior olfactory nucleus, both of which receive entorhinal input and provide feedback to the olfactory bulb, will require projection-specific manipulations combined with neural recordings. More generally, future experiments will be needed to determine how entorhinal cortex, piriform cortex, and AON interact to shape olfactory bulb representations during generalization.

Together, our findings identify an entorhinal feedback circuit that promotes successful generalization by suppressing neural representations of novel background stimuli. Activation of this circuit by an Ntsr1 agonist reduced abnormal sensory responses, improved neural coding, and restored behavioral performance in two genetically distinct mouse models of ASD. The ability of Ntsr1 agonists to engage this circuit following systemic administration, together with the defined anatomical site of action, provides a framework for pharmacological targeting of circuit-level mechanisms underlying sensory processing deficits in autism spectrum disorders.

## Supporting information

List of animals used

## Supplementary Material

### Systemic Ntsr1 activation increased performance on go trials

Novel background odors were presented together with either go target odors, for which mice were required to lick to receive a water reward, or no-go target odors, for which mice were required to withhold licking to avoid a timeout. In both *Cntnap2^−/−^* ^5^ and *Shank3B^+/−^* ^6^ mice without Ntsr1 agonist, the most common errors during novel background odor trials were misses, in which mice failed to lick in response to go target odors. In contrast, performance on no-go trials was relatively high because mice correctly withheld licking in response to no-go target odors in the presence of novel backgrounds.

We therefore asked whether the improvement in performance produced by the Ntsr1 agonist was driven by changes in go trials, no-go trials, or both. The improvement was primarily due to enhanced performance on go trials.

For *Cntnap2^−/−^* mice, performance on go trials increased from 39.2% (181 trials) to 74.4% (121 trials) (see **Supp. Figure 1A**). This increase was significant (p=1.5e-9, Fisher exact test) and was observed in all *Cntnap2^−/−^* mice. Similarly, for *Shank3B^+/−^* mice, performance on go trials increased from 52.3% (365 trials) to 87.6% (201 trials). This increase was significant (p=2.5e-18, Fisher exact test) and was observed in 5 of 6 *Shank3B^+/−^*mice (see **Supp. Figure 1B**).

The improvement in go-trial performance occurred while maintaining the already high performance on no-go trials. For *Cntnap2^−/−^*mice, no-go performance increased from 86.7% (181 trials) to 88.5% (139 trials), a difference that was not significant (p=0.73, single-tailed Fisher exact test) (see **Supp. Figure 1C**). For *Shank3B^+/−^* mice, no-go performance increased slightly from 71.5% (389 trials) to 75.6% (217 trials), which was also not significant (p=0.88, single-tailed Fisher exact test) (see **Supp. Figure 1D**).

Thus, the improvement in odor recognition produced by systemic Ntsr1 activation was primarily driven by increased responses to go target odors rather than changes in performance on no-go trials.

### Systemic Ntsr1 activation did not increase sniff rate in *Shank3B^+/−^* and *Cntnap2^−/−^* mice

Subcutaneous administration of the Ntsr1 agonist improved odor recognition performance for novel background odors in *Shank3B^+/−^* and *Cntnap2^−/−^* mice. Because increased sniff rates have been associated with increased task engagement, we asked whether the behavioral improvement could be explained by enhanced sniffing for novel background odors. However, neither basal sniff rate nor odor-evoked sniff rate was increased by Ntsr1 agonist administration.

Basal sniff rate was measured during the 2 s period preceding odor delivery. For *Cntnap2^−/−^* mice, the baseline sniff rate without Ntsr1 agonist was 2.13 sniffs/s with 0.95 CI (2.07–2.19), calculated over 1804 s. Following Ntsr1 agonist administration, the basal sniff rate decreased to 1.64 sniffs/s with 0.95 CI (1.58–1.70), calculated over a total period of 2628 s. The average basal sniff rate was reduced in 3 of 4 mice, although the change in the average sniff rate per animal was not significant (p=0.36, double-tailed t-test).

For *Shank3B^+/−^* mice, the baseline sniff rate without Ntsr1 agonist was 1.88 sniffs/s with 0.95 CI (1.84–1.92), calculated over 4412 s. Following Ntsr1 agonist administration, the basal sniff rate was 1.91 sniffs/s with 0.95 CI (1.86–1.96), calculated over 2772 s. The average basal sniff rate was reduced in 4 of 6 mice, and the difference in average sniff rate per animal was not significant (p=0.54, double-tailed t-test).

Odor-evoked sniff responses were also not increased by the Ntsr1 agonist. We analyzed sniff rates during the 750 ms period following target odor onset. During this interval, mice were exposed to both the novel background odor and the target odor that determined whether a lick response was required. To avoid potential confounds arising from differences in licking behavior between drug and control conditions, only correct trials were analyzed, and go and no-go trials were evaluated separately.

For *Cntnap2^−/−^* mice, the target-evoked sniff rate during go trials without Ntsr1 agonist was 3.54 sniffs/s with 0.95 CI (3.09–3.98), calculated over 69 presentations. Following Ntsr1 agonist administration, the sniff rate decreased to 3.02 sniffs/s with 0.95 CI (2.66–3.38), calculated over 90 presentations. The difference in the average response per mouse was significant (p=0.021, paired double-tailed t-test, n=4 mice).

The sniff response to no-go target odors was also reduced by the Ntsr1 agonist. The target-evoked sniff rate during no-go trials without Ntsr1 agonist was 3.84 sniffs/s with 0.95 CI (3.52– 4.15), calculated over 147 presentations. Following Ntsr1 agonist administration, the sniff rate decreased to 2.69 sniffs/s with 0.95 CI (2.40–2.98), calculated over 123 presentations. The difference in the average response per mouse was significant (p=0.0014, paired double-tailed t-test, n=4 mice).

For *Shank3B^+/−^* mice, the Ntsr1 agonist also did not increase odor-evoked sniff rates. During go trials, the target-evoked sniff rate without Ntsr1 agonist was 3.92 sniffs/s with 0.95 CI (3.64– 4.19), calculated over 191 presentations. Following Ntsr1 agonist administration, the sniff rate decreased to 3.39 sniffs/s with 0.95 CI (3.12–3.67), calculated over 174 presentations. The difference in the average response per mouse was not significant (p=0.34, paired double-tailed t-test, n=6 mice).

During no-go trials, the target-evoked sniff rate without Ntsr1 agonist was 3.19 sniffs/s with 0.95 CI (2.98–3.40), calculated over 278 presentations. Following Ntsr1 agonist administration, the sniff rate decreased slightly to 3.16 sniffs/s with 0.95 CI (2.88–3.43), calculated over 161 presentations. The difference in the average response per mouse was not significant (p=0.41, paired double-tailed t-test, n=6 mice).

Thus, the improvement in odor recognition produced by Ntsr1 agonist administration cannot be explained by increased sniffing behavior. Neither basal nor odor-evoked sniff rates were elevated by the drug, and several measures showed modest reductions in sniff rate despite improved behavioral performance.

### Ntsr1 is expressed in both medial and lateral entorhinal cortex, and systemic Ntsr1 activation increased c-Fos expression in both subdivisions

Within the entorhinal cortex, we compared Ntsr1 expression between the lateral and medial divisions. We analyzed three in situ hybridization experiments from the Allen Brain Atlas and quantified Ntsr1 expression energy in each subdivision. For each animal, Ntsr1 expression in the lateral and medial entorhinal cortex was normalized to total expression across the entire entorhinal cortex. The relative Ntsr1 expression was 51.2% ± 10.6% (mean ± s.e.m.) in the lateral entorhinal cortex and was nearly identical to the expression in the medial entorhinal cortex (48.8% ± 10.6%).

Ntsr1 agonist administration increased c-Fos expression in both subdivisions of the entorhinal cortex. In the lateral entorhinal cortex, c-Fos expression increased to 2.43 ± 0.71-fold relative to saline controls. In the medial entorhinal cortex, c-Fos expression increased to 1.32 ± 0.26-fold relative to saline controls.

### Long lasting behavioral improvement in *Shank3B^−/+^ and Cntnap2*^−/−^ mice after injection of Ntsr1 agonist in the entorhinal cortex

The infusion of the Ntsr1 agonist into the entorhinal cortex resulted in a longer-lasting improvement in odor recognition compared to delivery via the subcutaneous route. Following entorhinal injection in two *Shank3B*^+/−^ mice, elevated performance was observed on the day after injection, whereas after subcutaneous delivery, performance on odor recognition with novel background odors returned to pre-injection levels by the following day.

In two *Shank3B*^+/−^ mice, baseline performance before entorhinal infusion was 55.3% (n = 76 trials, 2 mice). Performance improved after injection of 4.2 nM PD149163 into the entorhinal cortex to 71.6% (n = 88 trials). The following day, performance remained high at 72.7% (n = 88 trials) and was not significantly different from performance on the day of injection (p > 0.99, Fisher exact test) [GO13.1]. Performance declined after 6 days to 60.2% (n = 88 trials), which was not significantly different from pre-injection performance (p = 0.53, Fisher exact test).

### Entorhinal injections of Ntsr1 agonist did not reduce correct responses on go trials in *Shank3B*^+/−^ and *Cntnap2*^−/−^ mice

Activation of Ntsr1 in the entorhinal cortex increased reaction times for novel background odors. We therefore asked whether the longer reaction times were accompanied by a reduction in licking responses to go target odors.

For *Shank3B*^+/−^ mice, performance on go trials increased from 64.2% (165 trials) to 77.0% (100 trials) following entorhinal delivery of the Ntsr1 agonist. This increase was significant (p = 0.039, Fisher exact test) and was observed in 3 of the 4 mice tested. For *Cntnap2*^−/−^ mice, performance on go trials decreased from 43.2% (111 trials) to 32.0% (50 trials) following entorhinal delivery of the Ntsr1 agonist; however, this decrease was not significant (p = 0.22, Fisher exact test).

Thus, the increase in reaction time produced by entorhinal Ntsr1 activation was not accompanied by a significant reduction in correct licking responses to go target odors.

In contrast, performance on no-go trials improved in both mouse models. For *Cntnap2*^−/−^ mice, performance on no-go trials increased from 71.2% (111 trials) to 94.8% (77 trials), a significant improvement (p = 2.57 × 10⁻, Fisher exact test). Similarly, for *Shank3B*^+/−^ mice, performance increased from 50.4% (133 trials) to 73.3% (120 trials), which was also significant (p = 1.86 × 10⁻, Fisher exact test).

Therefore, the increase in reaction time following entorhinal Ntsr1 activation cannot be explained by a generalized suppression of licking behavior, as correct go responses were not significantly reduced in either mouse model.

### Entorhinal injections of Ntsr1 agonist did not increase sniff rate in *Shank3B*^+/−^ and *Cntnap2*^−/−^ mice

Although systemic administration of the Ntsr1 agonist improved performance in novel background odor trials, this effect was not accompanied by an increase in sniffing frequency. We therefore asked whether the behavioral effects of direct entorhinal Ntsr1 activation could be explained by changes in odor sampling.

Neither basal nor odor-evoked sniff rates were increased following entorhinal delivery of the Ntsr1 agonist. Basal sniff rate was measured in a 2 s window preceding odor onset. In *Cntnap2*^−/−^ mice, basal sniff rate was 2.23 sniffs/s (95% CI: 2.15–2.31; 398 s total) without drug and 2.05 sniffs/s (95% CI: 1.95–2.15; 278 s total) following entorhinal Ntsr1 activation. The average basal sniff rate decreased in 2 of 3 mice, but this change was not significant (p = 0.74, two-tailed t-test). In *Shank3B*^+/−^ mice, basal sniff rate was 2.17 sniffs/s (95% CI: 2.10–2.25; 518 s total) without drug and 1.88 sniffs/s (95% CI: 1.81–1.95; 286 s total) with drug. The decrease was observed in all 4 mice but was not significant (p = 0.28, two-tailed t-test).

We next examined odor-evoked sniffing during the 750 ms following odor onset, separately for go and no-go trials.

In *Cntnap2*^−/−^ mice, evoked sniff rate during go trials was 3.89 sniffs/s (95% CI: 3.33–4.45; 48 presentations) without drug and 3.58 sniffs/s (95% CI: 2.72–4.64; 16 presentations) with drug (p = 0.71, paired t-test, n = 3 mice). During no-go trials, sniff rate was 4.27 sniffs/s (95% CI: 3.81– 4.73; 79 presentations) without drug and 3.98 sniffs/s (95% CI: 3.52–4.44; 73 presentations) with drug (p = 0.35, paired t-test, n = 4 mice) [GO17.1].

In *Shank3B*^+/−^ mice, evoked sniff rate during go trials was 6.03 sniffs/s (95% CI: 5.56–6.49; 106 presentations) without drug and 4.40 sniffs/s (95% CI: 3.93–4.87; 77 presentations) with drug (p = 0.014, paired t-test, n = 3 mice; one mouse did not contribute correct go trials under control conditions). During no-go trials, sniff rate was 5.41 sniffs/s (95% CI: 4.79–6.03; 54 presentations) without drug and 4.29 sniffs/s (95% CI: 3.86–4.72; 88 presentations) with drug (p = 0.38, paired t-test, n = 4 mice).

Across both genotypes and both trial types, entorhinal Ntsr1 activation did not increase basal or odor-evoked sniff rates. Thus, the improvement in odor-guided behavior cannot be explained by increased odor sampling.

## Methods

### Surgery

All surgical procedures were performed under ketamine/xylazine anesthesia, using aseptic technique and continuous monitoring of pedal withdrawal reflex and respiration. Adult mice (>60 days old, 20–25 g) were used for all procedures. Stereotaxic coordinates were based on the mouse brain atlas of Paxinos and Franklin. Mice were allowed to recover for 1 week before water restriction started. During imaging sessions, animals were head-fixed via the implanted titanium head-bar using a custom-built head-fixation apparatus.

### Head-bar implantation

Head-bar implantation was performed as previously described^5^. Mice were anesthetized with ketamine/xylazine (70/7 mg/kg, intraperitoneal) and maintained under anesthesia with supplemental doses as needed to diminish the pedal withdrawal reflex. Ophthalmic ointment was applied to prevent corneal drying.

The scalp was shaved and disinfected with betadine. Lidocaine and iodine were applied topically as local analgesic and antiseptic agents. Once a stable surgical plane was achieved, animals were secured in a stereotaxic frame using ear bars. A midline scalp incision (2–3 cm) was made to expose the skull. A titanium head-bar was affixed to the skull using light-cured dental cement (Vitrebond, 3M), positioned near the lambda suture to ensure stable head fixation for behavioral and imaging experiments.

### Cannula implantation in the entorhinal cortex

Mice undergoing entorhinal drug infusions were implanted bilaterally with guide cannulas during head-bar surgery. Cannulas were obtained from Protech International (formerly Plastics One) and consisted of a stainless-steel external guide cannula (26 gauge; 0.405 mm diameter; 5 mm tubing extending below the pedestal) and an internal injection cannula (33 gauge; 0.180 mm diameter), which extended 1 mm beyond the guide cannula tip during infusion.

Cannulas were targeted to the entorhinal cortex using coordinates 4.4 mm posterior to bregma and 3.0 mm lateral. The guide cannula was positioned to a depth of 1.5 mm below the brain surface to minimize tissue damage, while the internal cannula extended to 2.5 mm during infusions.

Initial pilot experiments in which the guide cannula was lowered to 2.5 mm resulted in entorhinal damage and impaired learning of a second odor pair at low concentration (250 ppm) without any backgrounds. To prevent this, implantation depth was reduced to 1.5 mm, and deeper targeting was achieved using the internal cannula.

Implantation was performed in a stepwise manner to minimize tissue disruption. Initial pilot experiments also show behavioral impairment in target recognition if the cannula lowering velocity was higher. The first 300 μm of descent was performed manually in a continuous motion (∼250 μm over 2 s) until brain surface contact was confirmed by visible dimpling. The next 700 μm was lowered in 100 μm increments with 2 min pauses between steps. The final 500 μm was lowered in 100 μm increments with 5 min pauses between steps to reduce tissue stress near the target region.

### Olfactory bulb imaging window

To measure olfactory bulb activity, *Cntnap2*^−/−^ × Thy1-GCaMP6f and *Shank3B*^+/−^ × Thy1-GCaMP6f mice were implanted with chronic optical windows over the olfactory bulb during the head bar and cannula surgery. The procedure followed previously established methods ^8^(Sturm et al., 2025), which modified an earlier skull-thinning approach (Otazu et al., 2015) to improve long-term stability.

The skull over the olfactory bulb was thinned using a dental drill until cortical vasculature became visible. A drop of Vetbond tissue adhesive was applied and allowed to polymerize for 5 min. Cyanoacrylate glue (Pacer Technologies, ZAP-A-GAP CA+) was then applied, and a 3.5-mm-diameter glass coverslip was pressed onto the skull and held in place for 10 min. The coverslip edges were sealed with dental acrylic.

### Drug preparation

PD149163 was purchased from Sigma (PD 149163 tetrahydrochloride hydrate, Sigma catalog: PZ0175). For subcutaneous injections, we dissolved PD149163 in saline (0.9% NaCl) at a concentration of 40 µg/ml (42.377µM). This dilution was delivered subcutaneously at a dose of 0.05 ml per 13 gr of mice, which corresponds to 0.13 mg/Kg. For direct injection into the entorhinal cortex, PD149163 was diluted in saline to a concentration of 4.2 nM. Injections into the entorhinal cortex were delivered into the entorhinal cortex at a volume of 1 µL, delivered over 5 minutes on each side of the entorhinal cortex. For two *Shank3B* mice, we used a lower concentration of 2.1 nM. With similar results as the 4.2 nM injection.

### Drug delivery

Subcutaneous injections of PD149163 were delivered before the behavioral session. Following the SQ injection, animals were returned to their cages for 30 minutes, before the start of the behavioral experiment. Animals that received the injections into the entorhinal cortex were first head fixed into the experimental apparatus. Internal cannulas were then inserted and the drug was injected. Behavioral experiment was started immediately following the delivery of the drug into the entorhinal cortex. For the cannula implanted animals, on the days without drugs we either injected them with saline into the entorhinal cortex (1 uL over 5 minutes) or did not receive any drug. To minimize the damage into the entorhinal produced by the infusion, we limited the sessions that mice received saline infusions.

### Behavioral training

Behavioral training was described before^5^ (Li et al., 2023). Mice were implanted with titanium head bars and water deprived for ≥7 days. Body weight was monitored daily and maintained at 80–85% of pre-deprivation weight by post-session water supplementation. Mice were trained in two stages: target-only training followed by target-in-known-background training. In target-only training, head-fixed mice learned go/no-go odor discrimination using two odor pairs (go: isopropyl butyrate, propyl butyrate; no-go: isobutyl propionate, ethyl propionate; 0.34% saturated vapor). Go responses were initially reinforced with water delivery (4 μL) and later required lick-based responses. No-go odors were introduced at low probability and progressively increased to 50% as performance stabilized. Odor concentrations were gradually reduced to 0.025%, and all analyses used sessions with 50% go probability. Time-outs were triggered by early licks or false alarms.

In target-in-known-background training, odor mixtures were introduced by pairing target odors with contextual backgrounds (0.025–0.1% saturated vapor). Background odor onset preceded target onset by 1.5 s, with (S)-(−)-limonene introduced 0.75 s after contextual background onset. Background odors persisted throughout the 3 s target period. Mice were trained over multiple days as background concentrations were increased stepwise to final levels (target 0.025%, background 0.1%).

### Sniff monitoring

Sniffing behavior was monitored using the same approach described by Li et al. (2023). Briefly, an airflow sensor (1000 SCCM AWM300V, Honeywell) was positioned opposite the animal’s nose, and airflow signals were low-pass filtered with cutoff frequency of 56.5 Hz to reduce high-frequency noise. Filtered signals were acquired at 1000 Hz using a NI USB-6003 data acquisition board.

### Whole-brain c-Fos quantification following systemic PD149163 administration

To determine the brain areas activated by systemic delivery of Ntsr1 agonist, we quantified c-Fos expression throughout the brain using tissue clearing, light-sheet imaging, and automated whole-brain cell counting. This approach provides an unbiased and quantitative measure of brain-wide neuronal activation following drug administration.

PD149163 was administered subcutaneously at a dose of 0.13 mg/kg, and c-Fos expression was compared with saline-injected controls. Experiments were performed in 3 *Shank3B*^+/−^ mice and 3 *Cntnap2*^−/−^ mice. Following injection, mice were returned to their home cages for 2 h before tissue collection.

Mice were transcardially perfused with saline followed by 4% paraformaldehyde (PFA). Brains were processed using the SHIELD protocol to preserve tissue architecture and protein antigenicity ^79^. Samples were post-fixed in SHIELD reagent and subsequently cleared for 7 days using Clear+ delipidation buffer. Cleared brains were labeled with rabbit anti-c-Fos antibody (3.5 μg per brain) and propidium iodide (24 μL per brain) as a nuclear counterstain.

Following refractive index matching, intact brains were imaged using a SmartSPIM light-sheet microscope (LifeCanvas Technologies) at 3.6× magnification, with a z-step of 4 μm and an xy pixel size of 1.8 μm. Three imaging channels were acquired: 488 nm autofluorescence for background subtraction, 561 nm propidium iodide, and 647 nm c-Fos immunofluorescence.

Image volumes were automatically registered to the Allen Mouse Brain Atlas using the propidium iodide channel. c-Fos-positive neurons were detected automatically, and the location of each cell was mapped onto the corresponding Allen Brain Atlas region. This procedure enabled quantification of the density of activated neurons within each brain structure across the entire brain. All tissue clearing, imaging, registration, and automated cell detection were performed by LifeCanvas Technologies.

### Analysis of spatial expression of Ntsr1

Regional expression of Ntsr1 was quantified using publicly available in situ hybridization datasets from the Allen Mouse Brain Atlas. Data from three independent Ntsr1 mRNA in situ hybridization experiments (Allen Brain Atlas experiment IDs 71325306, 73519704, and 80342232) were downloaded using the Allen Brain Atlas Application Programming Interface (API) through custom MATLAB scripts.

Gene expression levels were quantified using the Allen Brain Atlas expression energy metric. Expression energy is defined as the sum of expression pixel intensities divided by the total number of pixels within a given brain region, thereby normalizing expression measurements for regional volume. Expression energy values were extracted for the brain regions analyzed in this study and used to compare relative Ntsr1 expression across brain areas.

### Analysis of axonal projections from the entorhinal cortex

Axonal projections originating from the entorhinal cortex were quantified using publicly available anterograde tracing data from the Allen Mouse Brain Connectivity Atlas. The dataset consists of adult C57BL/6J mice that received stereotaxic injections of a Cre-independent recombinant adeno-associated viral tracer expressing enhanced green fluorescent protein (AAV2/1.pSynI.EGFP.WPRE.bGH) under the human synapsin I promoter, resulting in fluorescent labeling of neuronal cell bodies and their axonal projections throughout the brain.

We analyzed 11 experiments in which the viral injection site was located within the entorhinal cortex (Allen Brain Connectivity Atlas experiment IDs: 640081415, 642967852, 180297139, 272414403, 114472145, 126116848, 146553971, 585026021, 127139568, 113226232, and 112672974).

Brains were imaged using serial two-photon (STP) tomography and registered to the Allen Mouse Common Coordinate Framework. Projection strength to each target region was quantified using the Allen Brain Connectivity Atlas measurements of fluorescent axonal signal within anatomically defined brain structures. For each experiment, the fraction of total labeled axons projecting to each brain region was calculated by normalizing the projection signal in each target area by the total projection signal across all analyzed regions. These values were used to compare the relative strength of feedforward projections to the hippocampus and feedback projections to olfactory structures, including the piriform cortex, anterior olfactory nucleus, olfactory tubercle, and main olfactory bulb.

### Widefield calcium imaging of olfactory bulb output during entorhinal Ntsr1 activation

To determine how activation of Ntsr1-expressing neurons in the entorhinal cortex alters olfactory bulb output, we performed widefield calcium imaging in Thy1-GCaMP6f mice expressing GCaMP6f in mitral and tufted cells, the principal output neurons of the olfactory bulb^61^. Thy1-GCaMP6f mice were crossed with either *Cntnap2*^−/−^ or *Shank3B*^+/−^ mice as previously described (Sturm et al., 2025). Imaging was performed through the chronic olfactory bulb windows described above.

Imaging experiments were performed in 2 *Cntnap2*^−/−^ × Thy1-GCaMP6f mice and 2 *Shank3B*^+/−^ × Thy1-GCaMP6f mice. All mice were implanted with bilateral guide cannulas targeting the entorhinal cortex. During behavioral testing, PD149163 (4.2 nM, 1 μL per site) was infused bilaterally into the entorhinal cortex over 5 min. Imaging sessions obtained following PD149163 infusion were compared with sessions in which mice received either saline infusions or no infusion. A total of 20 imaging sessions were analyzed, including 5 sessions with PD149163 infusion and 15 sessions without PD149163 infusion.

The imaging setup was previously described in Li et al. (2023) and Sturm et al. (2025). Widefield fluorescence imaging of the olfactory bulb was performed using a pair of back-to-back SLR lenses, consisting of a 50 mm f/1.4 lens used as the objective and a Tamron AF 90 mm f/2.8 Di SP AF/MF 1:1 macro lens coupled to a scientific CMOS camera (CS2100M, Thorlabs). The camera was fitted with a long-pass emission filter with a cut-on wavelength of 500 nm (FELH0500, Thorlabs). This optical configuration provided a native spatial resolution of 3.3 μm per pixel. Images were acquired with 4 × 4 binning, resulting in a final resolution of 13.2 μm per pixel.

Images were acquired at 40 Hz. Excitation was provided by a 470 nm LED (M470L4, Thorlabs) equipped with a GFP excitation filter (MF469-35, Thorlabs) and a diffuser (ACL2520U-DG6-A, Thorlabs) to produce uniform, pattern-free illumination. Illumination intensity was adjusted to the minimum level that produced reliable odor-evoked fluorescence responses in order to minimize photobleaching. The objective aperture was maintained fully open (f/1.4) throughout imaging sessions to maximize light collection.

Calcium imaging data were analyzed in MATLAB (R2023b). Image registration across recording sessions was performed using the MATLAB function *imregconfig* with the “multimodal” configuration, aligning each session to a common reference image. For each pixel, a normalized fluorescence signal (ΔF/F₀) was computed, with F₀ defined as the mean fluorescence during the 7 s baseline period preceding odor onset.

To reduce low-frequency spatial components associated with global odor-evoked activity, ΔF/F₀ images were convolved with a Gaussian kernel (σ = 199 μm), and this low-pass filtered component was subtracted from the original signal. The resulting images were subsequently smoothed using an additional Gaussian convolution (σ = 39 μm) to reduce high-frequency spatial noise.

Regions of interest (ROIs) corresponding to putative glomeruli were manually identified using ImageJ ^80^. ROIs were drawn on activity maps generated by averaging responses across all presentations of identical odor combinations and were refined using ΔF/F₀ images from individual presentations of novel-background odor mixtures. An ROI was included in the analysis if it exhibited an average response greater than 1 z-score during the 500 ms window following target odor onset in at least one of the eight mixtures containing standard background odors. This image-processing pipeline followed previously established procedures for olfactory bulb calcium imaging analysis^8^.

For *Cntnap2*^−/−^ mice, 732 ROIs were analyzed in sessions without PD149163 (104.57 ± 22.25 ROIs per session, mean ± s.d.; 7 sessions) and 229 ROIs were analyzed following PD149163 infusion (114.50 ± 3.54 ROIs per session; 2 sessions). For *Shank3B*^+/−^ mice, 759 ROIs were analyzed in sessions without PD149163 (94.88 ± 36.79 ROIs per session; 8 sessions) and 264 ROIs were analyzed following PD149163 infusion (88.00 ± 19.70 ROIs per session; 3 sessions).

Because behavioral performance differed between drug and no-drug sessions, neural analyses were restricted to correct trials. For novel-background trials, 149 correct trials without PD149163 and 53 correct trials with PD149163 were analyzed in *Cntnap2*^−/−^ mice. In *Shank3B*^+/−^ mice, 93 correct trials without PD149163 and 209 correct trials with PD149163 were analyzed.

To quantify responses to background odors, average ROI activity was measured during the 500 ms window immediately preceding target odor onset, when only the familiar and novel background odors were present. To quantify responses during target presentation, average ROI activity was measured during a 400 ms window beginning 100 ms after target odor onset. Fluorescence responses were expressed as z-scored activity values.

To evaluate the consistency of drug effects across odors, average responses were calculated separately for each of the 11 novel background odors and compared between drug and no-drug conditions. Statistical significance was assessed using binomial tests. To account for differences between animals and genotypes, multivariate linear models were fit to individual ROI responses with PD149163 treatment as the primary factor and animal identity and genotype included as covariates. Statistical analyses were performed in MATLAB.

---

**Supplementary Figure 1.**
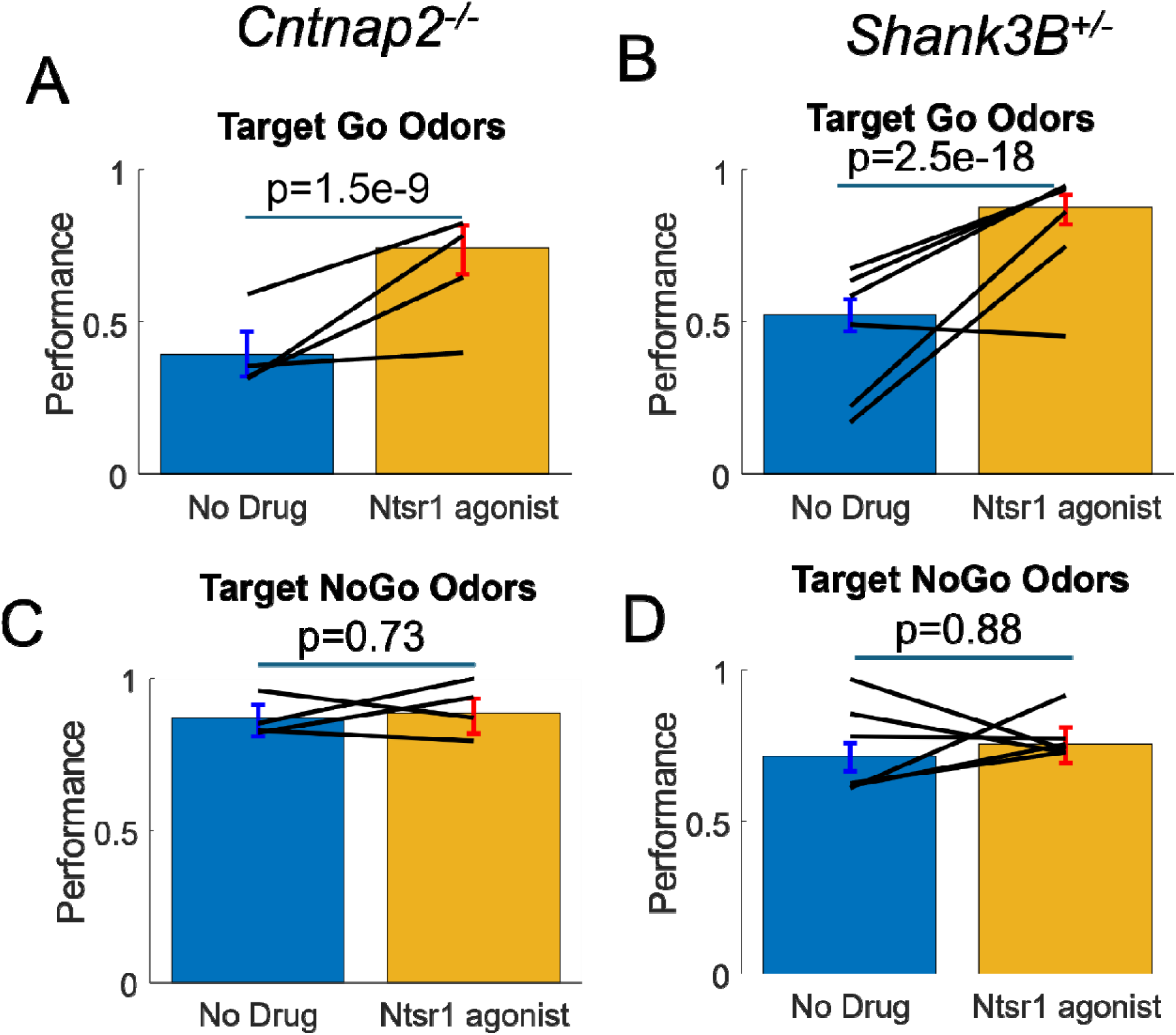
Systemic Ntsr1 activation improved performance on go trials. (A,B) Performance on go trials during novel-background testing in *Cntnap2*^−/−^ (A) and *Shank3B*^+/−^ (B) mice before and after subcutaneous administration of PD149163. Ntsr1 agonist significantly increased correct responses to go target odors in both mouse models. (C,D) Performance on no-go trials during novel-background testing in *Cntnap2*^−/−^ (C) and *Shank3B*^+/−^ (D) mice. Performance on no-go trials was not significantly altered by PD149163. Bars represent the mean, error bars indicate the 95% confidence interval, and black lines connect measurements from individual mice.

**Supplementary Figure 2.**
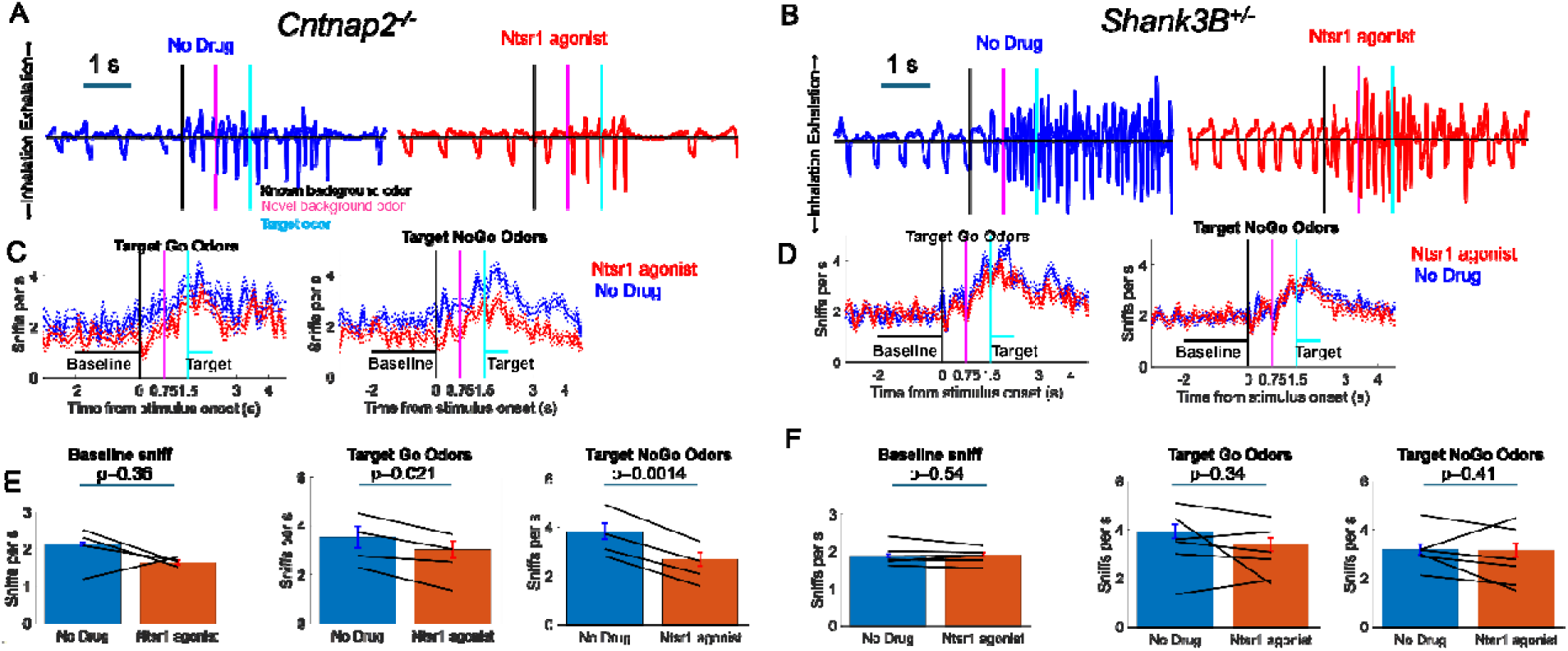
Systemic Ntsr1 activation did not increase sniff rate during novel-background odor presentation. (A,B) Representative respiration traces from a *Cntnap2*^−/−^ mouse (A) and a *Shank3B*^+/−^ mouse (B) during presentation of a novel background odor with and without subcutaneous administration of the Ntsr1 agonist. (C,D) Mean sniff rate traces from *Cntnap2*^−/−^ (C) and *Shank3B*^+/−^ (D) mice during novel-background odor presentation with and without Ntsr1 agonist administration. Shaded regions indicate the 95% confidence intervals obtained by fitting sniff rate events with a Poisson model. (E,F) Mean sniff rate with and without subcutaneous Ntsr1 agonist administration in *Cntnap2*^−/−^ (E) and *Shank3B*^+/−^ (F) mice. Ntsr1 activation did not significantly increase sniff rate in either genotype.

**Supplementary Figure 3.**
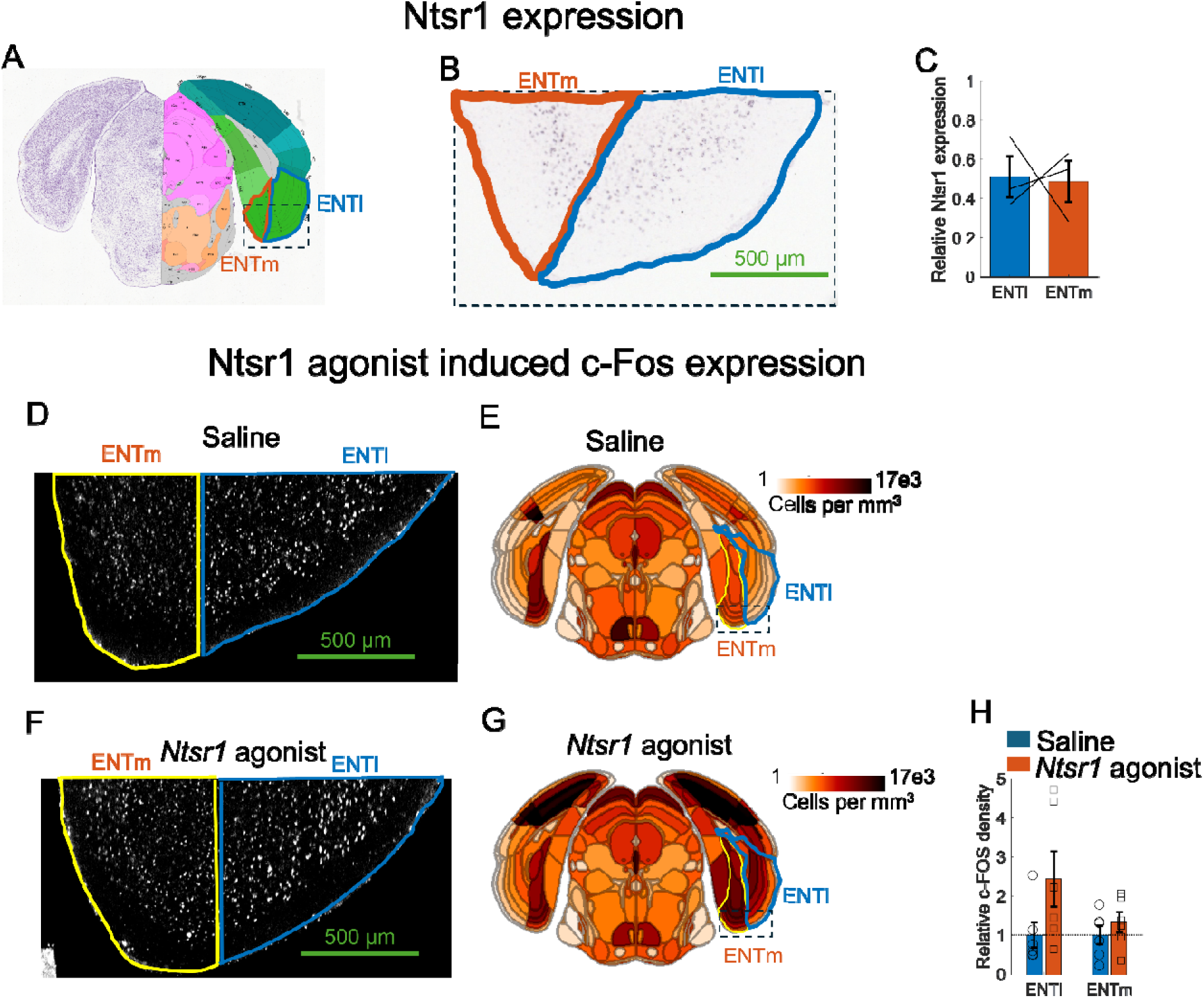
Ntsr1 is expressed in both medial and lateral entorhinal cortex, and systemic Ntsr1 activation increased c-Fos expression in both subdivisions. (A) Coronal atlas section showing the medial and lateral divisions of the entorhinal cortex. (B) Representative *in situ* hybridization section from the Allen Brain Atlas showing similar Ntsr1 mRNA expression in the medial and lateral entorhinal cortex. (C) Quantification of Ntsr1 expression from three Allen Brain Atlas datasets showing similar expression levels in the medial and lateral entorhinal cortex. (D,F) Representative sections from saline-treated and PD149163-treated mice showing increased c-Fos expression in both the medial and lateral entorhinal cortex following systemic Ntsr1 activation. (E,G) Average c-Fos density maps from saline-treated (n = 3) and PD149163-treated (n = 3) mice showing increased c-Fos expression in both entorhinal subdivisions. (H) Relative c-Fos density in the medial and lateral entorhinal cortex following systemic PD149163 administration. Bars represent the mean ± s.e.m.; symbols represent individual hemispheres.

**Supplementary Figure 4.**
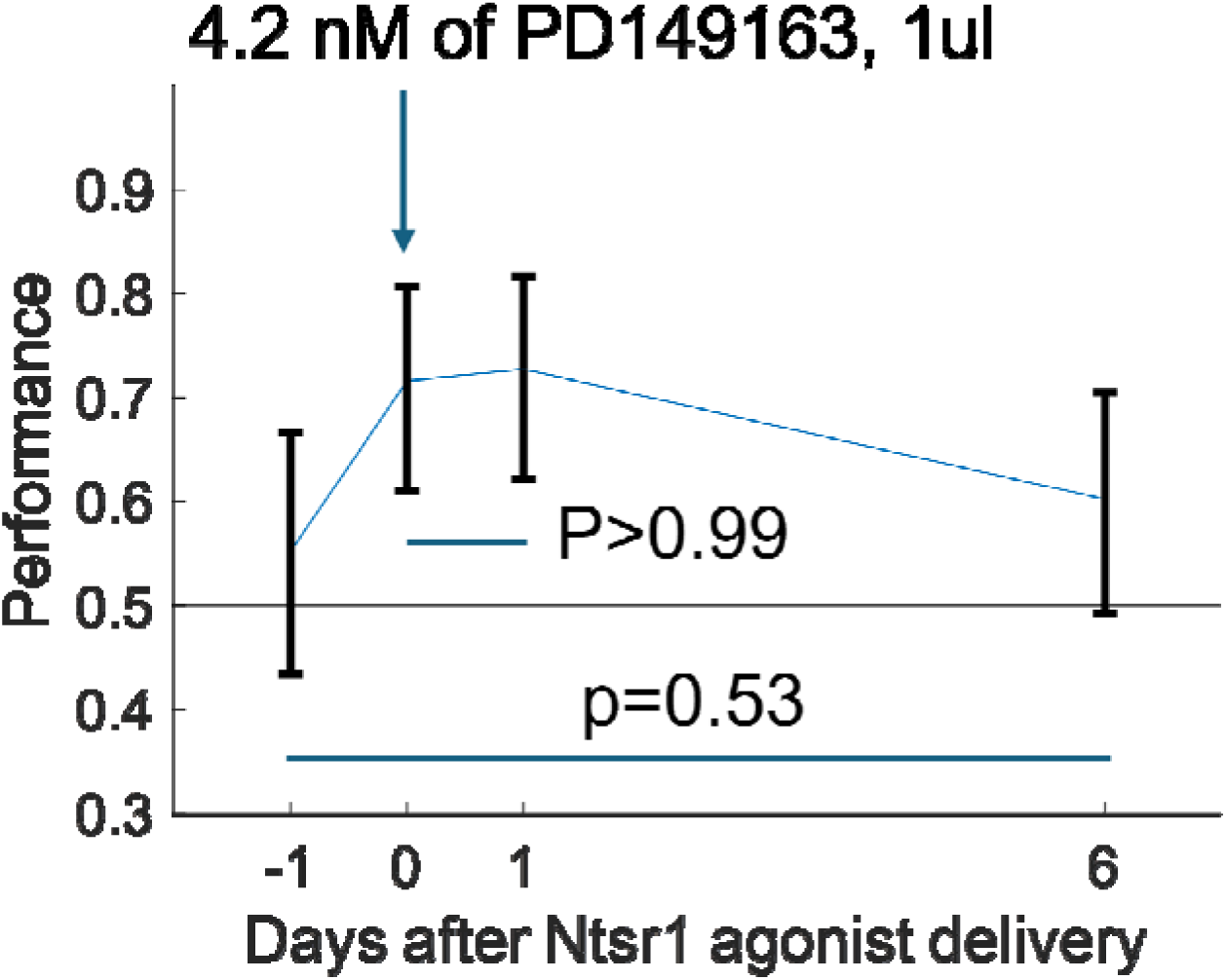
Behavioral improvement following entorhinal Ntsr1 activation persisted for at least six day. Performance of two *Shank3B*^+/−^ mice on novel-background odor trials before and after bilateral entorhinal infusion of 4.2 nM PD149163. Performance remained elevated on the day following infusion but returned to baseline 6 days later. Bars represent the mean and error bars indicate the 95% confidence interval.

**Supplementary Figure 5.**
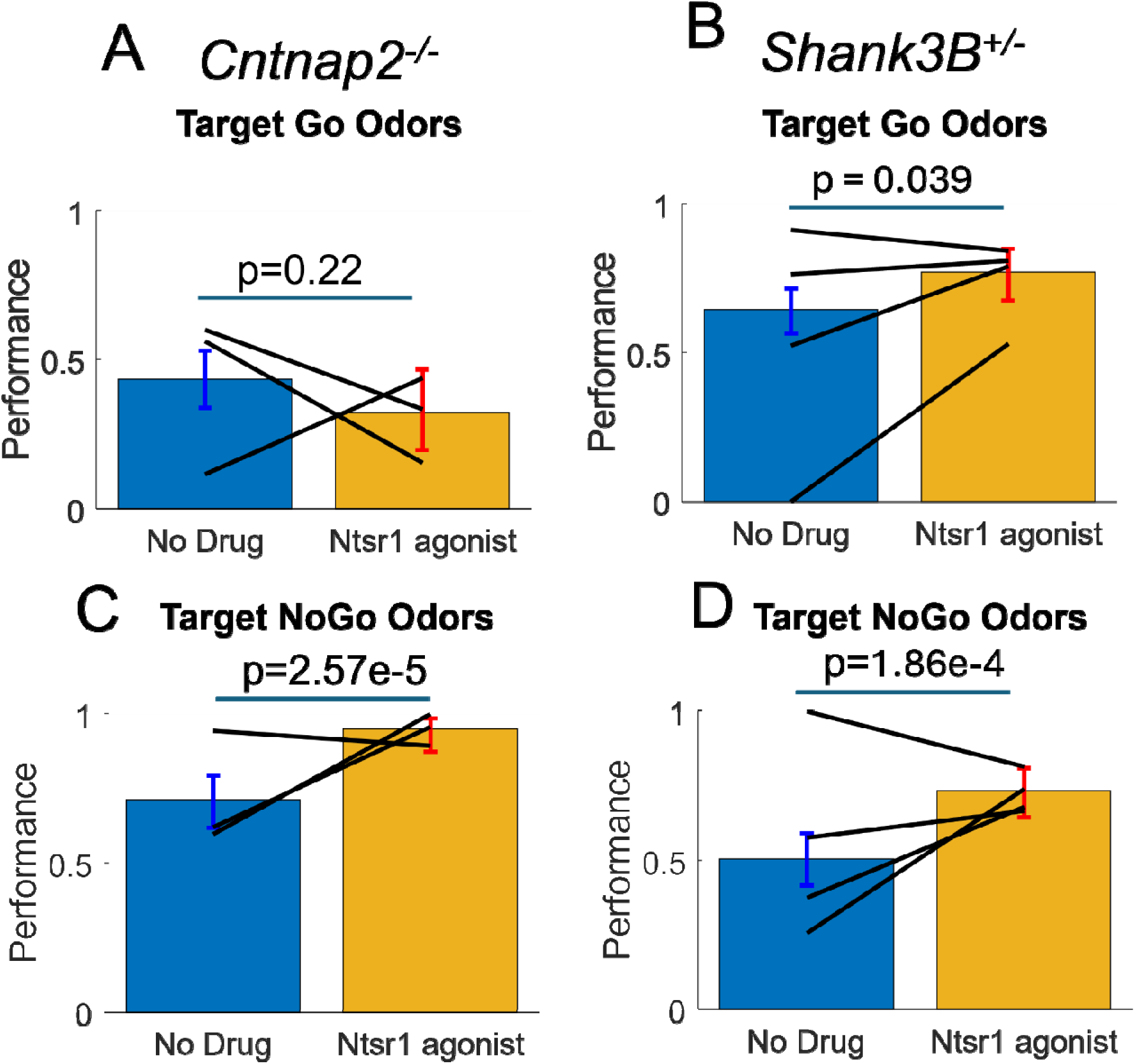
Entorhinal Ntsr1 activation did not impair go responses despite increasing reaction time. (A,B) Performance on go trials during novel-background testing in *Cntnap2*^−/−^ (A) and *Shank3B*^+/−^ (B) mice before and after entorhinal administration of PD149163. Correct responses to go target odors were not significantly reduced by entorhinal Ntsr1 activation. (C,D) Performance on no-go trials during novel-background testing in *Cntnap2*^−/−^ (C) and *Shank3B*^+/−^ (D) mice. Performance on no-go trials increased following entorhinal PD149163 administration. Bars represent the mean, error bars indicate the 95% confidence interval, and black lines connect measurements from individual mice.

**Supplementary Figure 6.**
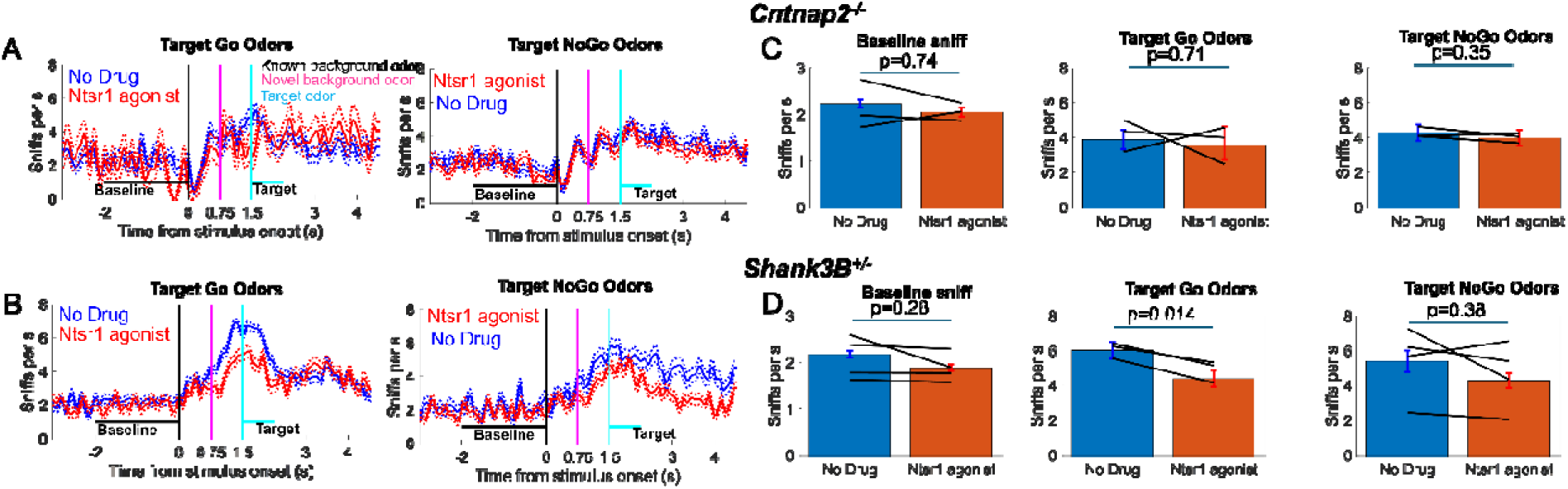
Entorhinal Ntsr1 activation did not increase sniff rate during novel-background odor presentation. (A,B) Mean sniff rate traces from *Cntnap2*^−/−^ (A) and *Shank3B*^+/−^ (B) mice during novel-background odor presentation with and without Ntsr1 agonist infusion into the entorhinal cortex. Shaded regions indicate the 95% confidence intervals obtained by fitting sniff rates with a Poisson model. (C,D) Mean sniff rate with and without Ntsr1 agonist infusion in *Cntnap2*^−/−^ (C) and *Shank3B*^+/−^ (D) mice. Ntsr1 activation did not significantly increase sniff rate in either genotype.

